# Single-molecule fluorescence microscopy demonstrates fast dynamics of the variant surface glycoprotein coat on living trypanosomes

**DOI:** 10.1101/2022.08.03.502583

**Authors:** Marie Schwebs, Torsten Paul, Marius Glogger, Leonard Forster, Brooke Morriswood, Jörg Teßmar, Jürgen Groll, Markus Engstler, Philip Kollmannsberger, Susanne Fenz

## Abstract

The fluidity of *Trypanosoma brucei*’s dense coat of GPI-anchored variant surface glycoproteins (VSGs) is fundamental for the survival of the parasite. In order to maintain the integrity of the coat, it is recycled on the time scale of a few minutes. This is surprisingly fast as endo- and exocytosis take place in the same small membrane invagination called the flagellar pocket. Here, we present measurements of VSG dynamics on the single-molecule level in living trypanosomes. A large number of short protein trajectories sampling the parasite’s surface were analysed in two distinct scenarios: diffusion and directed motion. To this end, we employed a previously published algorithm and implemented two extensions to consider rim effects as well as localisations errors inherent to single-mole tracking. Neglect of the latter can have a significant distortive effect on the measured diffusion coefficient; in our case resulting in an underestimation by 20 %. We found large heterogeneity in the local diffusion coefficients and velocities with a surprisingly high average value of 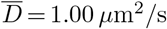 and 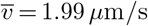, respectively. To decide on the locally dominant motion mode, we present a guideline based on random walk simulations. We find that VSG dynamics is indeed dominated by diffusion. Complementary simulations on long time scales not accessible in the experiment showed that passive VSG randomisation is fast enough to prevent re-endocytosis newly exocytosed VSGs and to accomplish turnover of the full VSG coat within a few minutes.

**Author summary:** Single-molecule tracking in biological systems often suffers from trajectories being too short to obtain statistically robust decisions on the present motion mode. We faced this issue when investigating the dynamics of the protein surface coat of African trypanosomes. To address the question whether diffusion or directed motion governs coat dynamics, we have adopted an algorithm based on temporal decomposition and spatial binning of an ensemble of single-molecule trajectories. We introduced several extensions to the original approach, including the consideration of localisation errors inherent to single-molecule tracking. This improvement alone already prevented the diffusion coefficient from being underestimated by 20 %. We analysed the coat dynamics in two scenarios, diffusion and directed motion, and offer a decision guideline to identify the locally dominating motion mode. Our extended algorithm is available to the scientific community via GitHub. For the trypanosome surface coat we found that the motion is indeed mainly characterised by diffusion with a surprisingly high diffusion coefficient. This finding solves a long-standing question how the parasite maintains its protein coat.

## Introduction

Extracellular pathogens of vertebrates are continuously exposed to their host’s immune response. This situation imposes a large evolutionary pressure on the pathogens to develop escape mechanisms. An outstanding example is the bloodstream form (BSF) of the unicellular eukaryotic parasite *Trypanosoma brucei*, the causative agent of human African trypanosomiasis (sleeping sickness) and nagana in livestock. Trypanosomes are fast-moving flagellates and their motility promotes dissemination within the host and thus, the establishment of an infection [1]. The parasite is covered with a dense monolayer of variant surface glycoproteins (VSGs). The fluidity of this surface coat is essential to escape the adaptive immune response of the host by antigenic variation [2, 3] and antibody clearance [4].

The homodimeric VSG is inserted into the outer leaflet of the plasma-membrane by a glycosylphosphatidylinositol (GPI) anchor at each C-terminus [5], which allows lateral mobility of the protein. Proper maintenance of the VSG coat’s integrity requires the VSGs to be shuffled through the flagellar pocket (FP), a plasma membrane invagination which accounts for only 5% of the total surface area and is the sole site of endo- and exocytosis. In 2004, the turnover of the total VSG surface pool was determined to be ^~^ 12 minutes which is surprisingly fast for a supposedly purely diffusive process [6]. A first attempt to measure VSG dynamics was undertaken already in the 1988 by Bülow et. al [7]. To date, studies addressing the dynamics of the VSG coat used fluorescence recovery after photobleaching (FRAP). This technique operates on a micrometer and second scale and allows to study the diffusion of a molecule ensemble. The measured diffusion coefficients were in the range of 0.01 - 0.03 *μ*m^2^/s [7–9] and were not sufficient to explain fast randomisation of VSG molecules and consequently fast turnover of a surface area of 100 *μ*m^2^. This raises the questions whether directed processes facilitate rapid VSG randomisation to prevent immediate re-endocytosis of exocytosed VSGs or to allow for fast turnover of the full coat. In the present study we address these questions using single-molecule fluorescence microscopy.

Swimming *T. brucei* can reach velocities of 42 *μ*m/s in the blood of their host [10]. This motility poses challenges for live-cell imaging. For the FRAP measurements, trypanosomes were embedded in either gelatine, restricting the acquisition to room temperature, or agarose gels, requiring additional interference with ATP household due to the insufficient stiffness of the hydrogel. Recently, an efficient, cyto-compatible immobilisation strategy was introduced that allows for immobilisation at the nanometer scale [11]. It relies on a cross-linkable hydrogel, which combines gel stiffness and cyto-compatibility, enabling us to image living trypanosomes at the physiological temperature of their host (37°C) without ATP depletion.

In this study, we characterised the dynamics of individual VSG molecules within the dense surface coat at high spatial and temporal resolution. Hereby, we checked the presence of directed motion towards the FP entrance. For this purpose, we established a cell line expressing a marker for the flagellar pocket entrance and labelled a small fraction of surface VSGs with an NHS-dye. Embedding trypanosomes in a cross-linkable hydrogel facilitated imaging under the physiological temperature of their host at 37°C. Due to the small size and large curvature of trypanosomes, the trajectories obtained were short and a reliable method for quantification was needed. Such a method was introduced by Hoze and Holcman [12]. Here, we adapted it to our data and extended it by two aspects: the consideration of the static and dynamic localisation error as well as a spatial filter for the correction of a projection error at the surface rim. This evaluation approach enabled to calculate maps of local diffusion coefficients and velocities, respectively. In addition, we offer a guiding principle for the decision whether diffusion or directed motion is predominant at a given site on the trypanosome map. Finally, we confirm by simulation that the measured diffusion coefficient signifies randomisation of VSG molecules fast enough to ensure escape of exocytosed VSGs from the pocket and efficient turnover of the full coat.

## Results

### Experimental layout

To investigate the diffusion properties of VSG molecules in the surface coat in relation to the entrance to the flagellar pocket (FP), we generated a BSF trypanosome cell line which expresses an N-terminally eYFP tagged TbMORN1. TbMORN1 is a main component of the hook complex [13, 14] which is wrapped around the FP neck as depicted in cyan in Fig 1 A. The hook complex was chosen as a reference structure due to its close proximity to the plasma membrane, while the pocket itself extends inside the trypanosome. Thus, the axial offset between the VSGs in the outer leaflet of the plasma membrane and the TbMORN1 molecules of the reference structure is reduced to a minimum. The new cell line was characterised with respect to cell growth and TbMORN1 expression (see S1 Appendix). We exploited the intrinsic photoswitching properties of eYFP at high illumination intensities to reconstruct a super-resolved image of TbMORN1. It exhibited the hook like shape known from immunofluorescence analysis (IFA) of the endogenous TbMORN1 (Fig 1 B, C).

**Fig 1.**
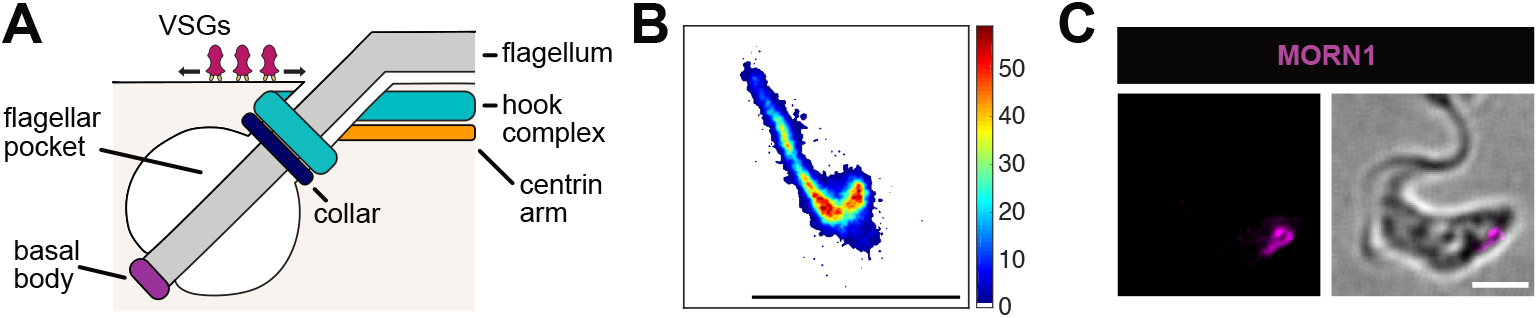
The hook complex of bloodstream form *T. brucei*. **(A)** The illustration depicts the position of the hook complex at flagellar pocket region. Scheme is not to scale. **(B)** Super-resolution image of the hook complex of a living, immobilised trypanosome expressing eYFP::TbMORN1. The quantity of single-molecule localisations is indicated by the colour code. The scale bar is 1.6 *μ*m. **(C)** Immunofluorescence analysis (IFA) of endogenous TbMORN1 in fixed trypanosomes. The endogenous protein was detected by using an antibody specific for TbMORN1 (magenta). The scale bar is 2 *μ*m.

To perform single-molecule measurements of VSG dynamics on living cells, the cells were first incubated with a nanomolar quantity of Atto-NHS 647N dye. This ensured a low label density suitable for. As 95% of the cell surface is covered by VSG molecules, there was a high probability that the emitters were labelled VSGs. Subsequently, the cells were embedded in a hydrogel of 10% (w/v) polyethylene glycol-vinylsulfone and 5.6% (w/v) thiol functionalised hyaluronic acid with a Youngs modulus of 105±29 kPa. Efficient nanometric immobilisation and survival of cells on the timescale of one hour were verified as in Glogger et. al [11]. Imaging was conducted at 100 Hz and at the hosts physiological temperature of 37°C. Subsequently, VSG trajectories were generated from the obtained VSG-Atto-NHS 647N localisations using a probability-based algorithm [15]. The dynamics of individual VSGs in the surface coat was investigated on 20 cells. Due to the small size of the parasite (length ~ 20 *μ*m, diameter ~ 4 *μ*m), was particularly challenging. As a consequence, the majority of the trajectories consisted of a length of less than 10 steps. However, 4.3 x 10^4^ ± 1.8 x 10^4^ trajectories were found on average per trypanosome resulting in sufficient sampling statistics to achieve reliable information on spatially resolved VSG dynamics.

### Computational details

For the analysis of data, the mean-squared displacement (*MSD*) is frequently used to derive a diffusion coefficient (*D*) which is defined for one dimension by Eq (1):

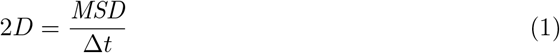

with Δ*t* describing the time lag. It is important to notice that in order to achieve statistically reliable results, the *MSD* may only be calculated for the first 10 - 20% of time lags of each trajectory of an ensemble, as the error in the *MSD* increases with increasing time lags [16]. Thus, long trajectories are required.

However, trajectories obtained from individual VSG emitters were short. In order to gain reliable information from these short trajectories and to evaluate trajectories also in the scenario of directed motion, we adapted the method introduced by Hoze and Holcman [17]. This method relies on a combination of temporal decomposition of trajectories and spatial binning of information to generate cell surface maps of diffusion and directed motion, respectively [12, 17]. Our adaptation of the algorithm included two extensions: The incorporation of the correction of localisation errors and the implementation of a spatial projection filter at the rim of the trypanosome surface projection (see S4 Appendix). Here, the adapted and extended algorithm is referred to as shortTrAn (short Trajectory Analysis) and its implementation is depicted in Fig 2.

**Fig 2.**
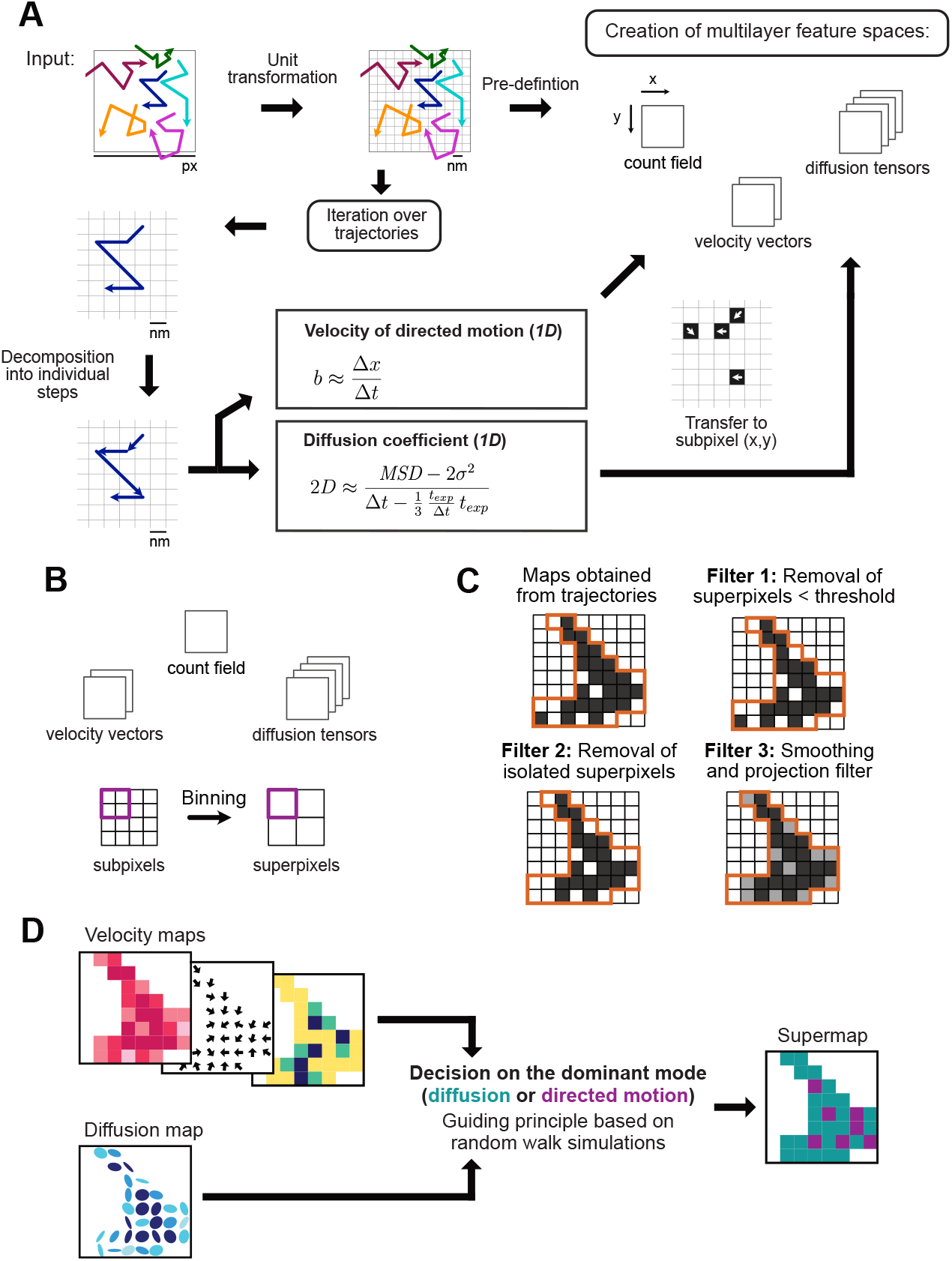
Workflow of the short trajectory analysis (shortTrAn). **(A)** The x- and y-positions of the trajectory steps served as input. First, the camera pixelation was transformed to a nanometre scale. The analysis iterated over each trajectory. Each trajectory was decomposed into its individual steps. Each step was evaluated for the distinct scenarios of diffusion and directed motion, respectively. Hereby, the velocity of a directed motion and the diffusion coefficient was calculated. The results were transferred to the subpixels of the steps origin in the pre-defined matrices, so-called feature spaces, which represented the image. The count field contained the number of steps originating from each subpixel. While the directed motion vector matrices comprised the velocities in x- or y-direction and the diffusion tensor matrices contained the diffusion coefficients of the xx-, xy-, yx- and yy-direction. **(B)** Binning subpixels to larger superpixels improved the statistics. **(C)** Application of spatial filters removed superpixels with low statistics and isolated superpixels. Finally, the data was smoothed out and a projection filter was applied. **(D)** The results for the diffusion and directed motion scenario were represented in respective maps. A decision guideline based on random walk simulation identified the locally dominant motion mode which were represented in a single supermap.

The localisation errors mentioned above are arising from sampling errors inherent to the single-molecule technique. These are referred to as the (i) static and (ii) dynamic localisation error [18]. Both errors influence the measured *MSD* and consequently *D* especially when it is calculated from *MSD* at Δ*t* =1. (i) The static localisation error depends on the localisation precision *σ*. The localisation precision of each emitter is mainly limited by the number of obtained photons and can be determined particularly well for an immobile emitter. In the community, this error was recognised almost two decades ago and is accounted for by

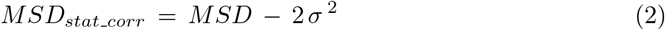

for one dimension [18–20]. The static localisation error thus adds a constant offset to the real *MSD*. This leads to an overestimation of the diffusion coefficient especially at small time lags or small diffusion coefficients if not taken into consideration properly (*D_stat_corr_* < *D*). (ii) The dynamic localisation error is caused by the exposure time required to collect enough photons from a single molecule. As a consequence, a moving emitter is imaged at positions averaged over the exposure time. The dynamic localisation error impacts the effective time point at which a molecule is localised and may be corrected for as follows:

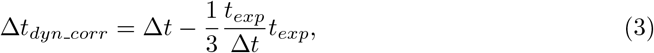

where *t_exp_* denotes the exposure time [18, 20–22]. This error, in contrast to the static localisation error, results in an underestimation of the diffusion coefficient, if it is not corrected for (*D_dyn_corr_* > *D*). Both errors have the largest effect on *MSD* (Δ*t* = 1) and *D* (Δ*t* = 1), respectively. In summary, the two errors have an inverse effect on the measured diffusion coefficient. It is noteworthy that the errors’ relative contributions are unique to individual experimental set-ups and must therefore be determined for each experiment.

We display the result of our measurements of VSG dynamics as diffusion and directed motion maps of each trypanosome surface (Fig 2 D). Additionally, we visualised the (an)isotropy of the diffusion as ellipses in the diffusion map and key figures of directed motion in three maps: a velocity heat map, a quiver plot illustrating the direction, and a relative standard error map. *A priori*, it was unknown whether diffusion or directed motion dominated locally. In Section “Guideline for the decision on the dominant motion mode”, we present a guiding principle for the decision. However, we started from the null hypothesis that the dynamics of the VSGs is a diffusion process.

### Evaluation in a diffusion scenario revealed inhomogeneous VSG diffusion in the trypanosome surface coat

Fig 3A depicts an exemplary ellipse plot visualising the displacements in light of a diffusion model. The colour code reflects the amplitude of the local diffusion coefficient obtained from the mean of both ellipse axes, while the shape of the ellipses allowed to distinguish between isotropic (circles) and anisotropic (ellipse) diffusion. The deviation from a circle and thus isotropic diffusion can be quantified by means of the eccentricity. The evaluation of all obtained datasets comprising 25,391 trajectories in total revealed that spatial heterogeneity of the diffusion coefficients was present (*N* = 20 cells). In the case of the exemplary cell, the local diffusion coefficient varied between 0.07 *μ*m^2^/s and 2.60 *μ*m^2^/s with a median of 0.81 *μ*m^2^/s (Fig 3 B). While the diffusion coefficient of the superpixels varied greatly, the average diffusion coefficient of all 20 cells was comparable and was on average 1.00±0.15 *μ*m^2^/s (Fig 3 B). For each cell, at least one well-defined domain with a low diffusion coefficient could be observed, but generally several of these were present (circled in Fig 3 A). Another prominent feature visible in the ellipse plot were membrane areas characterised by aligned elongated ellipses indicating biased VSG diffusion along the major axis of the ellipses (see arrows Fig 3 A). A correlation of the flagellum with areas of biased diffusion could be excluded by comparison of the regions in the corresponding transmitted light images and conducting VSG tracking on a cell line which has the flagellum fluorescently labelled (S1 Fig).The location of the entrance to the flagellar pocket was identified by the outline of the hook complex which is wrapped around the FP neck while the handle points towards the cell anterior. Near the flagellar pocket, the amplitude of the diffusion coefficients and the eccentricity of the ellipses were similar to those of the rest of the cell surface.

**Fig 3.**
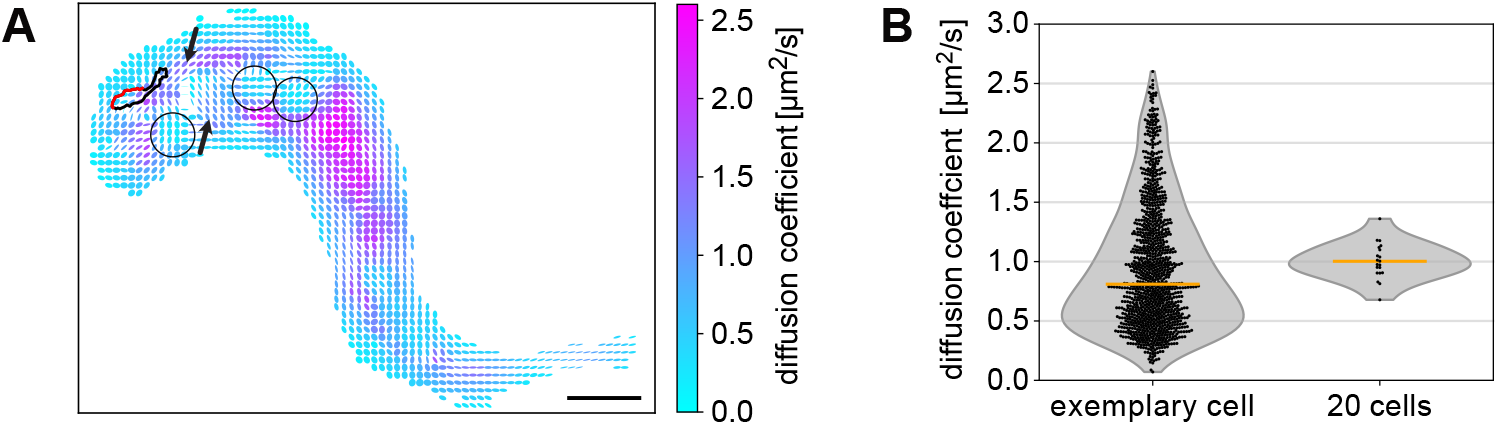
Diffusion scenario. **(A)** Local diffusion coefficients on the trypanosome surface. Diffusion information of superpixels are represented in ellipses.The amplitudes of the diffusion coefficients are colour-coded, while the shape of the ellipses encodes an- or isotropic diffusion behaviour. The outline of the hook complex is represented with a bold black line. The red highlighting indicates the area wrapped around the FP neck marking the entrance to the FP. Circles indicate domains of lower diffusion coefficients and arrows mark arrays of elongated ellipses. The scale bar is 1.6 *μ*m. **(B)** Distribution of the diffusion coefficients. The left box presents the spread of the local diffusion coefficients of the exemplary cell with a median of 0.81 *μ*m^2^/s (*N* = 1124 superpixels). The right box displays the distribution of average diffusion coefficients of all cells with a mean of 1.00 *μ*m^2^/s (*N* = 20 cells).

### Localisation errors have a strong effect on diffusion coefficients

As already mentioned, the technique itself suffers from sampling errors: The static and dynamic localisation error. The correction of both errors is particularly important in our case because the determination of the diffusion coefficient is based on the absolute values of the *MSD* (Δ*t* = 1). To determine the influence of both localisation errors on our measurements and to verify the necessity of the correction, we implemented the correction of the static error on the *MSD* and of the dynamic error on the time lag.

For the consideration of the static localisation error, the localisation precision σ must be ascertained. Three different approaches exist to obtain *σ*: (i) from the error associated with the determination of the x- and y-position by Gaussian fitting, (ii) by calculation of known parameters [23] or (iii) from the standard deviation of the localisations of an immobile emitter [24]. The results for *σ* by all three methods are displayed in S2 Appendix. The average *σ* of all three methods was 0.026 *μ*m. The correction of the dynamic localisation error was straightforward as all parameters were known *a priori*. The average two-dimensional diffusion coefficient corrected for both errors (*D_corr_*) over all 20 cells was 1.00 *μ*m^2^/s. In order to determine the net effect of both corrections contributing to the resulting diffusion coefficient, we back calculated an uncorrected diffusion coefficient for comparison:

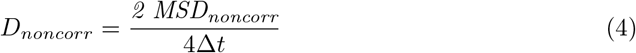

with

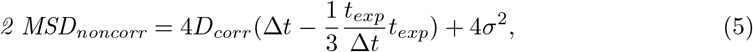

where *σ* = 0.026 *μ*m, *Deltat* = 0.01 s and *t_exp_* = 0.009 s.

We found a *D_noncorr_* of only 0.80 *μ*m^2^/s. Thus, if the static and dynamic localisation errors were ignored in the current experiment, the average real diffusion coefficient had been underestimated by 20 %. When we look at the relative contributions of both errors individually, we found that the overestimation of the static localisation error was minor (+ 8 %) in comparison to the underestimation by dynamic localisation error (−27 %). Even though, in principle, the localisation errors could cancel each other out, here we found a large net effect.

### Evaluation in a directed motion scenario showed locally high velocities of VSGs on the trypanosome cell surface

The results of the analysis of the local directed motion were depicted in three maps.

i. A velocity map which is a heat map representing the amplitude of the velocity (Fig 4 A). We observed a large variety of local velocities measured on the surface of a single cell. These ranged from 0.05 *μ*m/s to 10.35 *μ*m/s with a median of 1.75 *μ*m/s for the exemplary cell. The average over all 20 examined cells yielded 1.99±0.20 *μ*m/s (see also Fig 4 D).
ii. A quiver plot illustrated the direction of the motion (Fig 4 B). Interestingly, all cells exhibited one or several attractive foci marked by a set of arrows pointing to a common centre (circled in Fig 4 B). The attractive foci correlated with the domains of slow diffusion detected in Fig 4 A. Analysis of the dwelling time of particles in these superpixels by examining the cumulative fluorescence intensity indicated that these foci originate from confined particles. The corresponding fluorescence information is shown in S2 Fig. The confined particles were additionally very slow, which is reflected in the short length of the arrows (Fig 4 B) and by the small velocity at the centre of the sinks (circled in Fig 4 A). Again, near the flagellar pocket, neither the amplitude of the velocity nor the direction of the arrows encoding the directional information deviated from the rest of the cell surface area.

**Fig 4.**
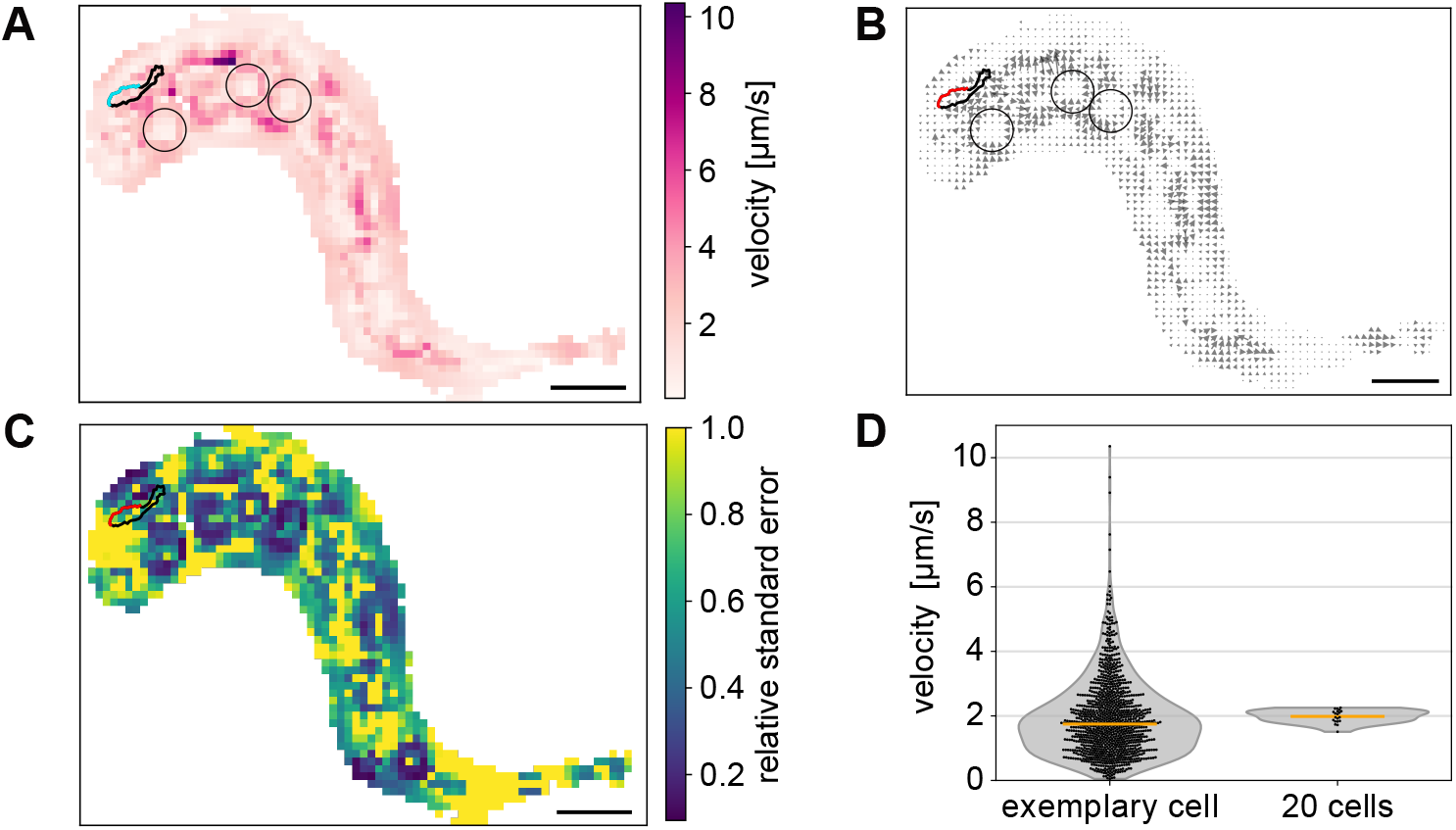
Directed motion scenario. **(A)** Local velocities on the trypanosome surface. The local velocities are colour-coded from white to purple. The bold black line represents the location of the hook complex, while the highlighting (cyan or red) marks the region wrapped around the flagellum marking the entrance to the FP. This is valid for (A), (B) and (C). Circles mark domains of low velocities. **(B)** Quiver plot depicting the directional information of the superpixel velocities. The arrow length correlates with the amplitude of the velocity. Circles mark attractive foci, characterised by arrows pointing to a common centre. **(C)** Heat map showing the local relative standard errors of the velocities. A threshold was applied which sets the relative standard error larger than one to the value one. The scale is 1.6 *μ*m. **(D)** Boxplot of the distribution of local velocities. The left box displays the spread of local velocities of the example heat map with their median of 1.75 *μ*m/s (*N* = 1124 superpixels). The right box presents the distribution of the average velocities from all cells (*N* = 20 cells). The mean value is 1.99 *μ*m/s and is indicated by the line. The exemplary cell is the same as in Fig 3.

In contrast to the calculation of the diffusion coefficients, the localisation errors have not yet been taken into account for the calculated velocities. (iii) A relative standard error map, which is a heat map, was generated to visualise the estimated error of the velocity due to localisation errors (see above). To consider the static localisation error, we acknowledged that for the measurement of the displacement and the associated velocity, the particle has to be localised twice. Each localisation was impaired by the limited localisation precision (*σ* = 26 nm). As a consequence, for a conservative estimate, the systematic error of the measured displacement in one dimension was ± 2*σ*. From this, we calculated the relative error of the mean velocity by classical error propagation. Finally, we arrived at the relative standard error of the mean velocity *SE_v_* by division by the square root of the number of displacements, *N*. We present the relative standard error, *SE_v_*, of each superpixel in a heat map (Fig 4 C). If we consider *N* = 1, an immobile particle could be wrongfully assigned a velocity of 5.2 *μ*m/s (*v*_0_ = 2*σ*/Δ*t* with Δ*t* = 10 ms). As a consequence, a true velocity for *N* = 1 has to be larger than 5.2 *μ*m/s. This threshold decreases with increasing statistics 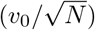. The larger the measured velocity became compared to the statistics dependent detection threshold, the smaller was the relative standard error. Consequently, the relative standard error allowed us to draw a conclusion about the reliability of the local velocity. We found that in all investigated cells, the *SE_v_* varied locally in the range of 0.029 to 215. For superpixels that had a relative standard error of the velocity equal or greater than 1, we concluded that there was no directed motion present or that it was below our detection limit. Thus, a cutoff at 1 was set for the representation of the relative standard error in the map (see Fig 4 C).

### Guideline for the decision on the dominant motion mode

In the evaluation of the VSG tracking data by shortTrAn, the same measured set of displacements was used to quantify VSG dynamics in the two distinct scenarios of diffusion and directed motion. However, *a priori* it is unknown which model is more suitable to quantify VSG dynamics locally. We were interested to identify superpixels where one type of motion clearly dominated aiming to resolve potential spatial segregation. Here, we present a decision guideline with the final results presented in supermaps. In these maps, each superpixel was colour-coded depending on the chosen mode. We suggest to use the eccentricity of the ellipses from the diffusion map as well as the relative standard error of the velocity as suitable parameter for the decision on the motion mode.

The first parameter, the eccentricity *ε* is a measure to determine the deviation of the diffusion tensor from a circular shape. The eccentricity is defined in a range *ε* ∈ [0,1], where the extrema 0 and 1 indicate a circle (diffusion) and line (directed motion), respectively. The second parameter, the relative standard error (*SE_v_*) equals 1, if the resulting velocity is solely caused by the inherent localisation uncertainty of the measurement. The relative standard error is very large either if the average velocity in a superpixel is very small or if the corresponding error is very large. The former will be true in case of slow directed motion or diffusion because random displacements add up to a zero net displacement. In contrast, fast directed motion will result in a very small relative error of the velocity (*SE_v_* → 0). In our measurements the eccentricity took all possible values between 0 and 1, whereas the relative standard error ranged between 0.029 and 215. Thus, we sought appropriate thresholds for both parameters to ascertain whether diffusion was present (*ε* < *ε_thres_*, *SE_v_* > *SE_v thres_*).

For this purpose, we simulated random walk processes. The 2D projection of the trypanosome surface is characterised by a large isoperimetric ratio of ~ 25*π*, which is only 4*π* for a circle of the same circumference. Thus, rim effects needed to be taken into consideration and simulations were conducted on a trypanosome-shaped mask. We simulated different datasets of trajectories with *D* = 1.0 *μ*m^2^/s from starting points randomly distributed on the trypanosome mask and analysed these datasets with shortTrAn. The number of data points and thus the statistics per superpixel were adjusted by the variation of the trajectory length (TL 15, TL50) or the number of trajectories within a dataset (2,000 - 10,000).

If we now look at the dependency of the eccentricity and the relative error on the statistics, we clearly see that there is not only one threshold value separating diffusion and directed motion, instead the threshold value must be chosen depending on the statistics (Fig 5 A, B). We set threshold values (*ε_thres_*, *SE_v thres_*) under the premises that 99% of the simulated diffusion data are assigned to diffusion and only 1% of the superpixels would be wrongly classified as non-diffusive (see Fig 5 C). In case of the eccentricity, a 99% threshold could be used as a value of 0 indicates isotropic diffusion. Thus, all superpixels with an eccentricity smaller than 0.79 (at *N* = 20, black arrow in Fig 5 C) were assigned to the diffusion mode. For the relative standard error, the threshold was set at 1% because a large relative error indicates diffusion whereas a relative error close to 0 signifies directed motion. Consequently, all superpixels with a relative standard error of larger than 0.45 (at *N* = 20, black arrow in Fig 5 C) were categorised in the diffusion group.

**Fig 5.**
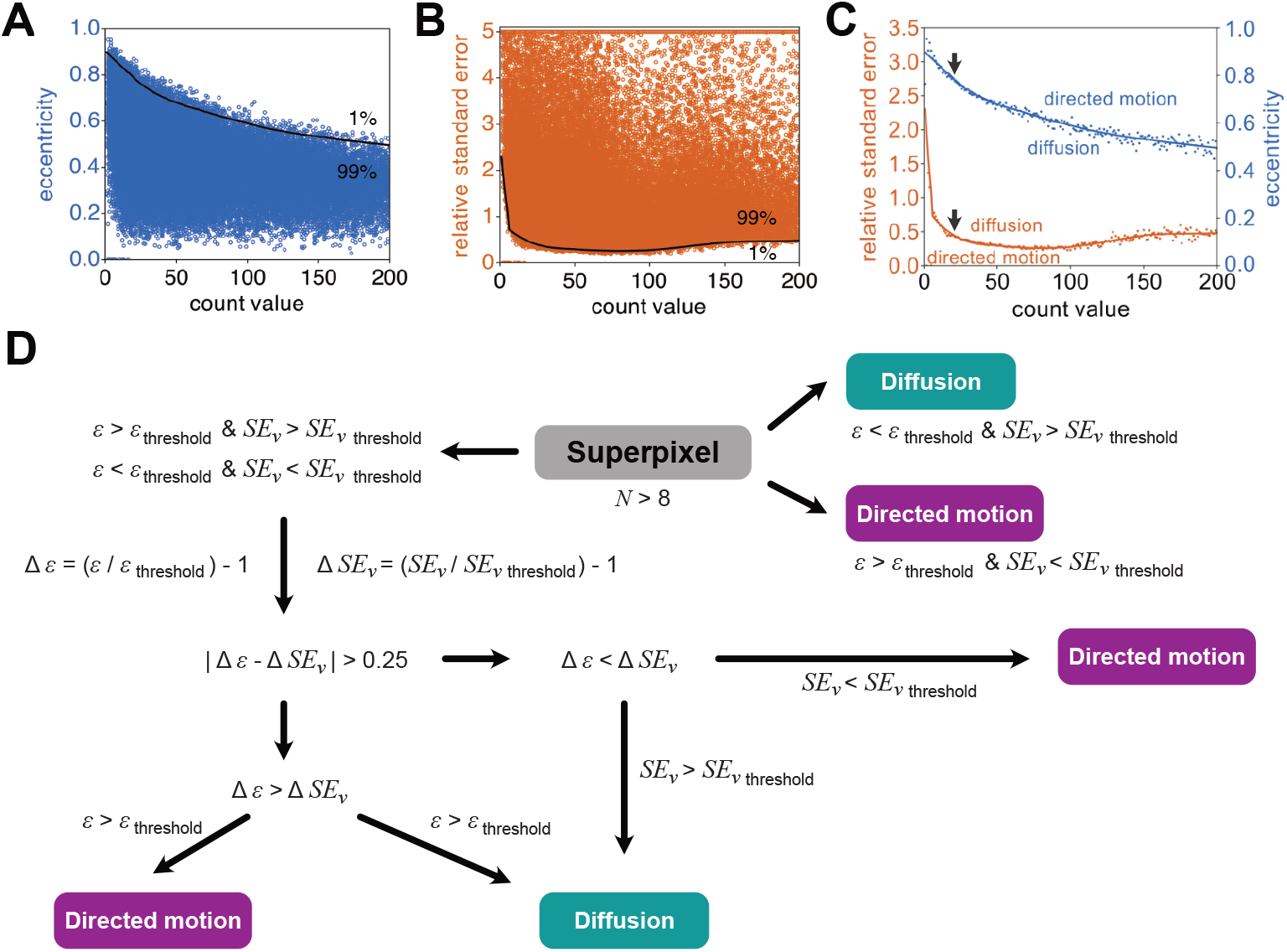
Decision guideline for the assignment of superpixels to one of the motion modes based on random walk simulations. **(A)** Dependency of the eccentricity extracted from the elliptic representation of the 2*D* diffusion tensors on the statistics, the count value of the corresponding superpixel. **(B)** Correlation of the relative standard error with the count value, the statistic of the corresponding superpixel. **(C)** Illustration of the thresholds set for the relative standard error and eccentricity to decide on the motion model. The threshold for the relative standard error was set to 1% of the data from (A) resulting in 99% of the data being assigned to diffusion. The eccentricity threshold was set to 99% of the data from (B) allowing 99% of the data to be attributed to diffusion. Black arrows indicate exemplary statistics at *N* = 20 mentioned in the text. **(D)** Flow chart illustrates the decision guideline using the introduced parameters and thresholds in (A), (B), and (C).

As expected, the eccentricity threshold decreased as the number of data points increased. However, even in case of *N* = 200, the eccentricity threshold was still 0.4 and thus above the theoretically predicted value of 0 indicative of perfectly isotropic diffusion. As anticipated, the threshold for the relative standard error of a hypothetical velocity was large at small statistics. As the number of data points per superpixel increased, two opposing effects took place. The standard error in the nominator decreased, while the average velocity in the denominator tended to zero 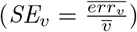. As a result the threshold of the relative standard error of the velocity first decreased strongly as *N* increased, reached a saddle point around *N* = 50 and finally increased again slowly before it saturated around *N* = 200. At low statistics (*N* < 8), the eccentricity was close to 1 and the relative standard error diverged. Thus, the unreliable regime was excluded from the analysis.

The superpixels were assigned according to the following scheme, which is also summarised in Fig 5 D. First, all superpixels were collected for which both parameters, eccentricity and relative standard error, pointed towards the same motion mode (follow the arrow to the right in the flow chart in Fig 5 D). If the measured eccentricity *ε* was small (*ε* < *ε_thres_*) and at the same time the measured relative standard error was large (*SE_v_* > *SE_v thres_*) the superpixel was assigned to the diffusion mode. Alternatively, if the attribution was *vice versa* the superpixel was assigned to the directed motion model. Afterwards, a large number of superpixels remained that could not be assigned as clearly because the two criteria were contradictory. In order to decide which of the criteria was stronger for a given case, we quantified how well the measured parameter could be distinguished from its corresponding threshold: Δ*ε* = (*ε*/*ε_thres_*) – 1 and Δ*SE_v_* = (*SE_v_/SE_v thres_*) – 1 (follow the arrow pointing downwards in the flow chart in Fig 5 D). Subsequently, the criterion that could be assigned with higher certainty to a motion model was identified by determining the difference between the two deviations from their threshold values (|Δ*ε* – *ΔSE_v_*|). If the difference between the two deviations was < 0.25, the superpixels remained unassigned because neither dominated the other. Conversely, if the difference was > 0.25, the criterion with the more significant difference to its threshold was used. If the eccentricity was found to be the parameter for the reliable assignment (Δ*ε* > Δ*SE_v_*), the superpixel was allocated to the diffusion in the case of *ε* < *ε_thres_* and to the directional model in the case of *ε* > *ε_thres_*. However, if the decisive parameter was the relative standard error (Δ*ε* < *ΔSE_v_*), the superpixel was assigned to diffusion if *SE_v_* > *SE_v thres_* or to directed motion if *SE_v_* < *SE_v thres_*.

Even after this refinement, unassigned superpixels remained. We categorised these orphan pixels with the help of their neighbours by using a moving box kernel (kernel size = 3 superpixels) to consult also superpixels on the diagonal. If the environment was uniform (7 and 8 superpixels of the same motion type), the centre was filled accordingly. This filtering process ran until no orphan superpixels could be filled by means of this principle.

In summary, the colour-coded supermaps allowed to grasp the predominant VSG dynamics on the trypanosome surface, with green superpixels indicating that the dominant motion mode was diffusion, magenta superpixels directed motion, and grey superpixels denoting orphan pixels that could not be assigned unequivocally.

### VSG dynamics are mainly characterised by diffusion, but round and elongated traps exist

Application of the guiding principle to the preceding results obtained from the shortTrAn evaluation of the VSG dynamics on 20 trypanosomes revealed a cell-to-cell variation in the relative prevalence of the locally predominating motion mode as well as in their spatial occurrence. Nevertheless, the supermaps could be consolidated into three different groups based on visual assessment of prominent features. Group 1 was characterised by a high proportion of diffusive superpixels and only a few, isolated superpixels assigned to directed motion (*N* = 7 cells). Group 2 was distinctive as the cells already had a larger proportion of superpixels assigned to directed motion, yet distributed as patches on the surface (*N* = 7 cells). Group 3 was characterised by a high proportion of directed motion-associated superpixels which clustered in contiguous, elongated arrays (*N* = 6 cells). Notably, the direction of the motion did not run along these arrays but pointed vertically into the centre, reminiscent of elongated traps. In most cases, there was even an area in the centre that was characterised by diffusion. The attractive foci found in the directed motion maps, correlating with slow diffusion, were still recognisable in the supermaps of all groups. They were characterised by a centre of diffusion circled by superpixels assigned to directed motion. From the directed motion maps we know that the direction of the motion pointed to the centre, which is reminiscent of a round trap. Representative supermaps of each group are shown in Fig 6A - C, while all 20 supermaps are displayed in S3 Fig.

**Fig 6.**
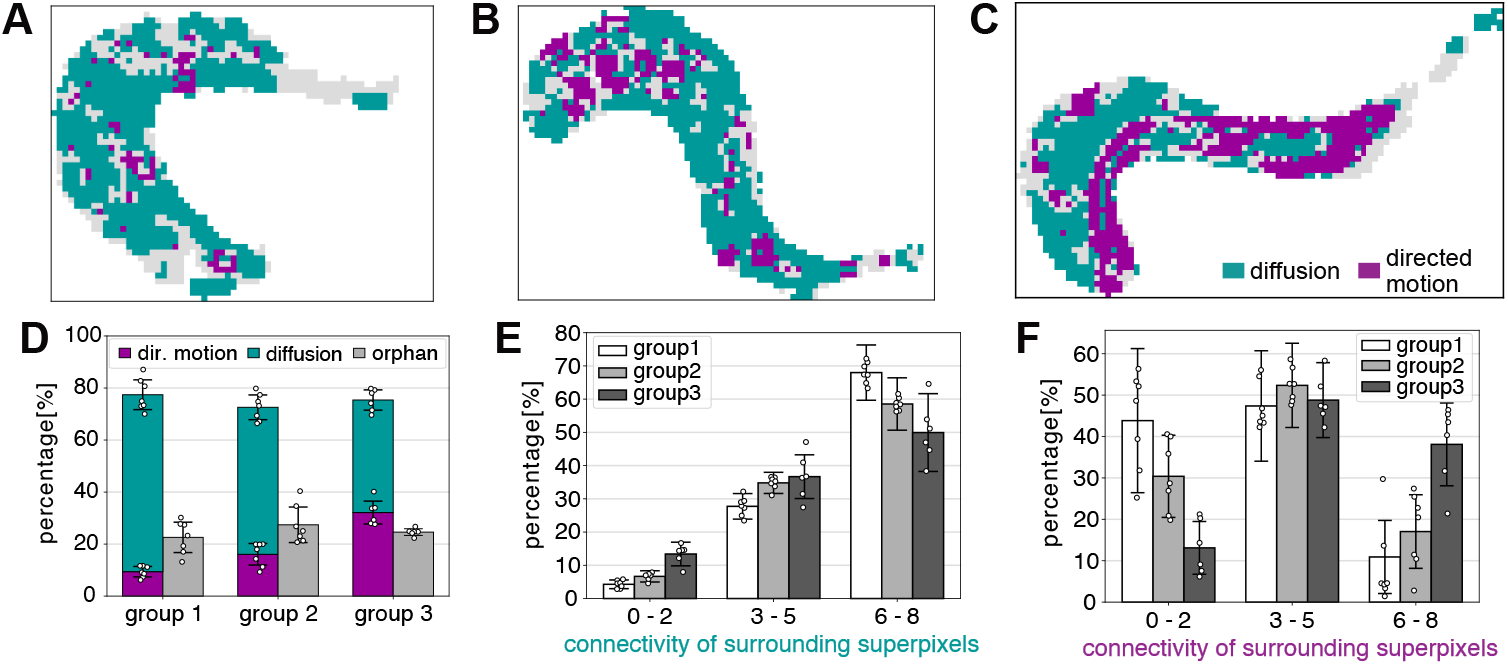
Exemplary supermaps and their quantification. **(A)** - **(C)** Supermaps of representative trypanosome surfaces. From left to right: group 1, group 2, and group 3. Diffusion is depicted in green, directed motion in magenta and orphan pixels in gray. One superpixel is of the size of 0.16 x 0.16 *μ*m. **(D)** Relative fraction of superpixels assigned to diffusion, directed motion or without assignment (orphan) within the supermap groups 1 - 3. Bars represent the mean percentage and error bars indicate the SD. Individual data points are shown as circles (*N* = 7 for group 1 and 2, *N* = 6 for group 3). **(E)** Percentage of superpixels assigned to diffusion with 0 - 2, 2 - 5, 6 - 8 neighbours exhibiting diffusive behaviour for the three different groups (group 1 in white, group 2 in gray, and group 3 in dark grey). Error bars represent the SD. Circles indicate the individual values of the supermaps within a group (*N* = 7 for group 1 and 2, *N* = 6 for group 3). **(F)** Percentage of superpixels attributed to directed motion with 0 - 2, 2 - 5, 6 - 8 neighbours exhibiting directed motion behaviour for the three different groups (group 1 in white, group 2 in grey, and group 3 in dark grey). The SD is shown by error bars. The individual values of the supermaps within the groups are depicted by circles (*N* = 7 for group 1 and 2, *N* = 6 for group 3).

Further quantification of the supermaps was performed by analysing the relative prevalence of the two motion modes as well as the connectivity of a superpixel’s associated motion model to the immediate surrounding. The relative prevalence in all three groups showed that more superpixels were assigned to the diffusion model (group 1: 68.0±5.7 %; group 2: 56.6±4.8 %; group 3: 43.3±3.9 %) over the directed motion model (group 1: 9.4±2.0 %; group 2: 16.1±4.1 %; group 3: 32.1±4.4 %) which is shown in Fig 6 D. While the fraction of orphan pixels was similar in all three groups, significantly more superpixels exhibited diffusion in groups 1 and 2, whereas diffusion and directed motion were almost equally prevalent in group 3. To quantify the connectivity of one type of motion, we extracted the number of the adjacent superpixels assigned to the same motion model. If up to two adjacent superpixels had the same motion model, they were grouped in category I. Category II and category III contained superpixels with three to five or six to eight neighbours of the same motion model, respectively. If the connectivity of pixels assigned to diffusion was examined (Fig 6 E), we found that 68% of all superpixels of cells in group 1 had six to eight neighbours assigned to the diffusion model (category III). If, on the other hand, we inspected the connectivity of superpixels assigned to directed motion (Fig 6 F), we found the highest percentage (38 %) of contiguous pixel (category III) in group 3. Additionally, most superpixels characterised by directed motion in group 3 had three to five neighbours assigned to directed motion indicating rather an elongated arrangement in one dimension than a arrangement in two dimensions. Together, this reflected the presence of elongated traps characterised by superpixels assigned to directed motion. In summary, quantification of the relative prevalence and the connectivity confirmed grouping by visual inspection and furthermore demonstrated that diffusion is indeed the dominating mode of VSGs in the surface coat.

### Simulations indicate that the measured average diffusion coefficient is sufficient to allow for fast VSG coat turnover

We checked whether the newly determined average diffusion coefficient of 1.00 *μ*m^2^/s was sufficient for a passive randomisation enabling the surprisingly fast turnover rate, implying that the entire VSG surface pool is internalised and recycled once in ~ 12 min [6]. The uptake through the small FP resembles the narrow escape problem (NEP) [25]. This problem deals with a Brownian particle which is confined to a bounded domain (cell surface), except for a small window (FP) through which it can escape. This process is characterised by the mean first passage time (*τ*).

First, we extracted *τ* from simulations of VSG diffusion on three trypanosome-shaped masks derived from the shortTrAn. The masks consisted of two layers to mimic the *3D* surface area of a trypanosome which is known to be 100 *μ*m^2^ [8]. Three trypanosome-shaped masks were selected which had an area of ~ 28 - 40 *μ*m^2^ per layer (see Table 1). The area per layer was still less than half of the total surface area due to our wide-field measurements that captured only the part of the membrane in focus. In accordance with the requirements for the NEP, the cells that served as blueprints for the masks were in the G1- or S-phase of the cell cycle and thus had only one FP. Only one layer contained the exit site, the FP region. The exit site region was extracted from the part of the hook complex wrapped around the FP entrance. We simulated 2,000 trajectories of diffusing particles per mask, with 1,996 - 2,000 trajectories escaping by hitting the FP exit site within a predefined time window of 500 s. The simulations revealed a *τ_sim_* of 50 - 78 s (see Table 1). We then complemented the determination of *τ_sim_* by theoretical considerations. As the VSG proteins are only diffusing in the plane of the membrane and thus effectively in two dimensions, we chose a theoretical model of a 2*D* domain with an escape site in the centre to calculate *τ_th_* according to the simplified Eq 6 [26]:

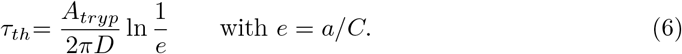

Here *A_tryp_* denoted the surface area of a trypanosome, *D* was the diffusion coefficient, *a* was the FP radius, and *C* was the circumference of the trypanosome mask. The calculation of *τ_th_* required the FP radius which was derived from a built-in function in MATLAB of the FP mask assuming a circular shape. *τ_th_* for the three cell masks was 51 - 73 s (see Table 1) and thus in good agreement with *τ_sim_*.

**Table 1.**
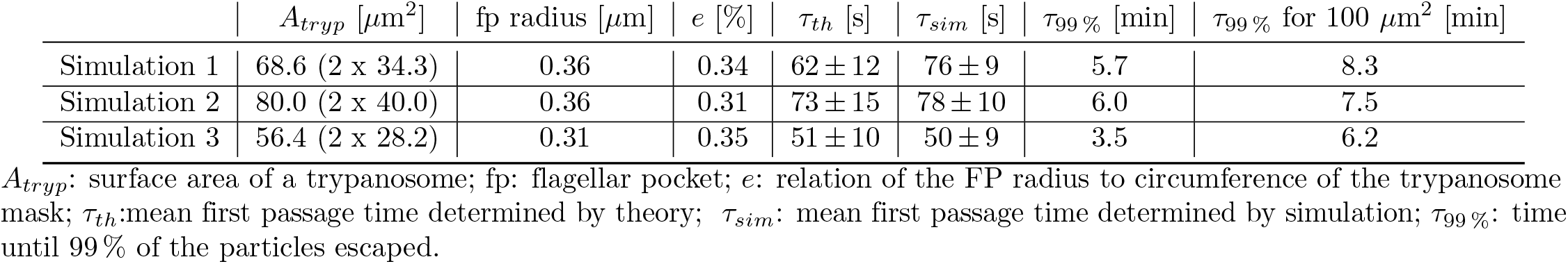
Determination of the mean first passage times (*τ*) of diffusive particles on trypanosome-shaped masks by theory and simulation.

Furthermore, to determine the time of VSGs distributed on the entire surface escaping through the FP, the time until all simulated VSGs have escaped through the FP was estimated. To this end, we calculated the time, *τ*_99%_, until 99% of the simulated trajectories escaped through the FP from the cumulative distribution of the individual escape times. *τ*_99%_ was found to be in a range of 3.5 - 6.0 min. As the used surface masks were smaller than 100 *μ*m^2^, we extrapolated *τ*_99%_ for all three simulations to 100 *μ*m^2^ based on the direct proportionality of the surface area (*A*) and the escape time (*τ*) in theory (see Eq 6). This finally led to an increase in *τ*_99%_ to 6.2 - 8.3 min, which emphasises that the randomisation of VSGs in the surface coat and thus a turnover time of ~ 12 min could be achieved by diffusion with an average diffusion coefficient of 1.00 *μ*m^2^/s.

## Discussion

In this study, we present single-molecule measurements of the dynamics of the surface coat proteins (VSGs) on living, but immobilised trypanosomes. Moreover, we introduce an algorithm for the analysis of short trajectories obtained from single-molecule fluorescence tracking (SMFT) taking into account the experimental localisation errors. We found a spatially inhomogeneous distribution of local VSG dynamics on the single-cell level. We identified diffusion as the overall dominant motion mode. Using the measured average diffusion coefficient in experimentally inspired simulations, we could demonstrate fast randomisation of VSGs enabling a turnover time of the total VSG coat within ~ 12 min [6].

In previous studies, VSG dynamics in the VSG coat of living trypanosomes were monitored with FRAP (fluorescence recovery after photobleaching) and was characterised with a diffusion coefficient in the range of 0.01 - 0.03 *μ*m^2^/s [7–9]. Here, VSG dynamics were investigated by SMFT which revealed a two orders magnitude higher diffusion coefficient of 1.00 *μ*m^2^/s.

We suspect that a combination of several aspects is at the origin of this large disagreement including differences due to the (i) imaging technique, (ii) labelling strategy as well as (iii) temperature at which measurements were conducted.

i. Although FRAP and SMFT are both techniques to monitor dynamics, they probe different spatial and temporal scales. SMFT operates at a nanometer and millisecond scale, while basic FRAP works in the micrometer and seconds range [27]. Furthermore, FRAP implicitly assumes that the specimen under investigation is homogeneous and that all particles of an ensemble behave similarly. However, the plasma membrane of a living cell is usually heterogenous (reviewed in [28]). As a consequence, FRAP yields an average of all possible distinct spatial behaviours as well as usually only one measurement per cell. SMFT, in contrast, distinguishes between individual proteins as well as provides spatially super-resolved information. Furthermore, the bleaching step in widefield-FRAP could contribute to the difference in diffusion coefficients gained by the two methods. The bleaching step requires high illumination intensities. As a consequence, the fluorescent label of the proteins anchored to the outer leaflet of the plasma membrane is affected on the top and bottom of the cell. This reduces the fluorescence emitting VSG pool for the recovery from outside the focus distorting the measurements and hence the results.
ii. Both techniques demand different labelling strategies. FRAP measurements require extensive labelling of the full surface coat to monitor the dynamics of the entire ensemble of surface proteins. Due to the high protein density in the VSG surface coat this could lead to artefacts due to steric hindrance, introduced by dye-dye interactions. SMFT has the advantage of working with nanomolar quantities to track individual emitters, which minimises the chance of steric hindrance. Moreover, the use of high dye concentrations, as in FRAP, will most likely not only result in labelling of the easily accessible VSGs, but additionally of less abundant surface proteins. Thus, the average dynamics of several, also much slower, protein families will be probed.
iii. In addition to the difference between the chosen imaging techniques, the environmental conditions also have an influence on the measured diffusion coefficient. Hartel and his colleagues carried out their measurements at room temperature, a limitation imposed by the chosen embedding strategy employing gelatine and agarose gels [8, 9]. However, the physiological temperature of the host of bloodstream form trypanosomes is 37°C and the diffusion coefficient itself depends on the temperature. This is described by the Einstein relation: *D* = *μ k_b_ T*, where *D* is the diffusion coefficient, *μ* is the mobility of the diffusive particle, *k_b_* is the Boltzmann’s constant, and *T* the absolute temperature. The temperature effect could account for an increase in the diffusion coefficient of ~ 6 %. Moreover, the fluidity of the lipid membrane is also influenced by the temperature directly via its phase behaviour. Usually, higher temperatures increase the fluidity of the lipid matrix and thus indirectly affect the dynamics of the anchored proteins. Bülow et. al performed FRAP at 37°C [7] and determined a diffusion coefficient similar to the ones determined by Hartel and his colleagues [8, 9]. However, they additionally had to interfered with the global ATP household to completely immobilise the parasites. A side-effect on VSG dynamics could not be excluded. In contrast, our work used trypanosomes embedded in thermostable hydrogels which allowed for measurements of the dynamics at 37°C without any interference in the ATP household.

It is noteworthy, the diffusion coefficient determined here is in better agreement with the diffusion coefficient reported for other GPI-anchored proteins in living cells [29, 30].

The evaluation in a diffusion scenario revealed local heterogeneities in the diffusion coefficients of VSGs on the trypanosome surface. Investigations of the VSG structure by X-ray scattering implies that they can adopt two main conformation [31]. The authors hypothesise that both conformations are an adaptation to varying VSG densities and obstacles in the surface coat. Furthermore, a correlation of the conformations with the amplitude of the diffusion coefficient is postulated [31].

Our approach of interpreting the single-molecule trajectories in two distinct scenarios, diffusion and directed motion, required a decision on the predominant motion mode at each location on the trypanosome surface. To this end, we combined the results of both scenarios pixel-wise in cell-specific supermaps by applying our decision guideline showed a substantial variation from cell to cell. Nevertheless, the cells could be categorised in three main groups, which were characterised by different proportions of superpixels assigned to diffusion or directed motion.

Interestingly, all three groups showed round domains in which superpixels of directed motion enclosed superpixels of diffusion in the centre. The movement of the directed motion pointed to the centre and reminiscent of round traps. We identified slow, confined emitters as the origin. We suggest four hypothesis for the observed confinement.

i. Labelled VSGs were endocytosed and located to endosomal structures. Their signal might be detected even though they are not in the plane of the plasma membrane because a widefield setup was used. In trypanosomes, the endosomal structures are localised between the FP and the nucleus. In our experiment, the round traps were also observed at the anterior part of the cell. Additionally, the round traps span at least three superpixels, corresponding to a diameter of 480nm. Thus, they are significantly larger than the average diameter of endosomes (*D*_endosomes_ = 135nm, [32]).
ii. NHS-dyes bind non-specifically to primary amines and consequently could have labelled not only VSGs but additionally some of the less abundant types of proteins. A suitable candidate could be a protein of the invariant surface glycoprotein (ISG) family. As some ISGs, like ISG-65 and ISG-75, are of a similar size to VSGs [33], the VSG coat might not shield this proteins completely. ISGs are 50-times less abundant than VSGs [33]. By using nanomolar quantities of the NHS-dye, the probability of labelling a rare ISG was reduced. This is in agreement with the low number of round traps and confined particles that were observed. Moreover, ISGs are transmembrane proteins [33, 34] and thus more likely to experience confined diffusion than GPI-anchored VSGs. The dynamics of ISG in the trypanosome membrane has not yet been addressed.

Another explanation for the observed confinement could be a physical confinement either (iii) due to interactions of the GPI-anchors with structures in the membrane influenced by the cytoskeleton or (iv) due to direct interaction of the VSGs with structural proteins. (iii) Raghupathy et. al reported that long chain GPI-anchored proteins couple to complementary inner leaflet nanodomains of phosphatidylserine which were associated with connector proteins of the actin cytoskeleton in Chinese hamster ovary cells [35]. In the case of trypanosomes, VSGs posses a short GPI-anchor and phosphatidylserine additionally represents only a very small fraction of the total cell lipidome in bloodstream form trypanosomes [36]. Furthermore, an association with lipid domains would be contradictory, as it would interfere with the functionality of the coat. For these reasons, it is considered unlikely that VSGs are trapped due to interleaflet coupling by their GPI-anchor. (iv) VSGs could interact with proteins present in the outer leaflet of the plasma membrane. Possible candidates are integral proteins which could also serve as connectors anchoring the microtubule cytoskeleton to the plasma membrane. In summary, all these hypothesis need to be addressed with further experiments.

To address the question why we observe three groups of VSG dynamics supermaps, we checked whether the cells in the groups are in fact in different cell cycle stages as the cells used for the experiment were not synchronised. However, a correlation of the groups with the proliferation stage of the cells could not be established, as dividing and non-dividing cells were present in each group.

One of the three groups introduced above revealed a unique characteristic: a high proportion of directed motion-dominated superpixels clustered in contiguous, elongated arrays. Notably, the direction of the particle movement did not run along these arrays but pointed vertically to the central line similar to the round traps. Again, there was an area in the centre characterised by diffusion. Therefore, these were identified as elongated traps. The obvious candidate imposing such a structure on the trypanosome membrane would be the flagellum. However, no spatial correlation could be found. Nevertheless, we assume that a passive effect is at the origin of the elongated traps, e.g. due to trapping by a structure.

Furthermore, we were interested in whether the newly determined and significantly higher diffusion coefficient would be sufficient to allow for a turnover of the VSG surface pool within ~ 12 min [6]. To this end, we performed elaborated experimentally inspired random walk simulations and conservatively estimated the time required for almost all VSGs of the surface coat to escape into the FP. These simulations emphasise that VSGs distributed on the entire surface area can reach the FP by free diffusion 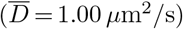 to accomplish a turnover time of ~ 12 min.

It was one aim of this study, to elucidate how it is ensured that exocytosed VSGs are not immediately re-endocytosed. To this end, we made the following estimation to compare the measured diffusion coefficient to the rate of endocytosed membrane area in trypanosomes. It is known that BSF trypanosomes exocytose 7 vesicles per second [6]. The average diameter of clathrin-coated vesicles is 135nm [6, 32]. Consequently, the endocytosed area was derived from the surface area of seven vesicles. The area endocytosed per second was calculated to be 0.401 *μ*m^2^. VSGs diffusing with 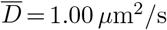 have a high probability to escape the endocytosed surface area, whereas VSGs diffusing with 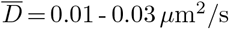 as determined by FRAP have a very small chance to reach the trypanosome surface. This implies that the fast diffusion coefficient alone is sufficient to distribute the majority of exocytosed VSGs to the cell surface and directing processes are not necessarily required.

To this point, an impact of the intrinsic motility of the trypanosomes on the VSG dynamics remained unconsidered. The influence of the forward motility of swimming trypanosomes on VSG dynamics cannot yet be investigated experimentally on a high-resolution scale as long acquisition times are required. However, it can be assumed that the additional energy introduced from the intrinsic motility will lead to intensified random mixing of laterally moving membrane-associated particles such as the VSGs. Thus, we can not exclude an influence on the VSG dynamics.

In summary, we presented measurements of VSG dynamics on the single-molecule level in living trypanosomes at the physiological temperature of their host. The adaptation of an evaluation algorithm for short trajectories and its improvement considering the static as well as the dynamic localisation error offers a standardised tool for the determination of reliable local diffusion coefficients and velocities. Furthermore, we provided a guideline to decide which of the motion modes is dominant on the nanometre scale. This algorithm together with the immobilisation of the parasites in a thermostable hydrogel provides an excellent tool to study the dynamics of surface proteins in trypanosomes. This tool might be also relevant for other protists such as *Leishmania*. The algorithm for shortTrAn as well as for the motion decision is now accessible to the research community via GitHub.

## Materials and methods

### Cell lines

A Lister 427, 13-90 cell line was established in which one allele of TbMORN1 (accession number Tb427.6.4670) was endogenously tagged with an eYFP at the N-terminus (eYFP:: MORN1). For this, the targeting construct contained the TbMORN1 5’-untranslated region (5’-UTR), blasticidin resistance gene, tubulin intergenic region, the eYFP coding sequence and parts of the TbMORN1 open reading frame (ORF) in a pCR4Blunt-TOPO cloning vector [13, 14]. Digestion with Cail and NdeI excised the targeting region for the transfection into bloodstream form trypanosomes. Only one allele was tagged to ensure detection of single molecules.

### Cell culture

The bloodstream form cells were cultivated in HMI-9 medium supplemented with 10% fetal calf serum at 37°C and 5% CO_2_. The integrated copies of pLew13 and pLew90 encoding the T7 polymerase and the tetracycline repressor were selected with 2.5 *μ*g ml^-1^ G418, 5 *μ*g ml^-1^ hygromycin, whereas the transgene eYFP::MORN1 was selected with 5 *μ*g ml^-1^ blasticidin. The cell density was kept below 5 x 10^5^ cells ml^-1^ to ensure exponential growth.

### Western Blot analysis

Western blotting was performed according to standard protocols. Trypanosomes were lysed in SDS loading buffer and ~ 1.4 x 10^6^ parasites were loaded per lane. Lysates were separated using SDS-PAGE, and the proteins were then transferred to a PVDF membrane. The detection of TbMORN1 was accomplished by using a rabbit anti-TbMORN1 antibody at 1:2,000. The eYFP-tagged fusion protein was detected by a rabbit anti-GFP antibody (1:5,000). As a loading control the histone variant H3 was detected with a guinea pig anti-H3 antibody (1:1,000, provided by Christian Janzen, University of Würzburg). Antibodies were used in combination with IRDye680 - or IRDye800 - conjugated secondary antibodies at 1:20,000 (Invitrogen) to allow signal detection with an Odyssey infrared scanner (LI-COR).

### Immunofluorescence analysis (IFA)

1×10^7^ trypanosomes were harvested by centrifugation (1400 xg, 10 min, RT) and resuspended in 1 ml trypanosome dilution buffer (TDB; 20mM Na_2_HPO_4_, 2mM NaH_2_PO_4_, 5mM KCl, 80mM NaCl, 1mM MgSO_4_, 20mM glucose). Cells were fixed in 2% paraformaldehyde (PFA) supplemented with 350mM NaCl for 15 min at RT. Fixed cells were washed three times with TDB and resuspended in 500 *μ*l TDB. Trypanosomes were transferred to poly-L-lysine-coated slides and allowed to settle for 30 min at RT. Cells were permeabilised with 0.5% Triton X100 in PBS for 15 min at RT. After washing once with PBS cells were blocked with 3% BSA in PBS for 30 min at 37°C. TbMORN1 was detected by a rabbit anti-TbMORN1 antibody diluted in 0.1% BSA in PBS (1:5,000, 1 h, RT). Subsequently, three washing steps with PBS were performed. For detection an Alexa594-conjugated secondary anti-rabbit antibody (Thermo Fisher Scientific) was used (1:2,000, 30 min, RT). After three washing steps with PBS and rinsing once with ddH_2_O, cells were mounted with Vectashield (Vecta Laboratories Inc.). Imaging was performed with an inverted-widefield fluorescence microscope (DMI600B, Leica) equipped with an immersion objective (100x, NA 1.40, OIL) and a DFC365FX camera (Leica) using the Leica Application Suite X software. Image processing was carried out with ImageJ.

### Preparation of glass coverslips

Thickness-corrected coverslips (22 x 22 mm, 17 *μ*m, Karl Hecht GmbH & Co KG) were sonicated in 2% Hellmanex for 10 min and extensively rinsed with deionised water. This process was repeated with deionised water before the coverslips were dried in the oven at 80°C and treated with plasma using ambient air for 30 min (PDC-002 CE, Harrick Plasma). The coverslips were stored in deionised water and used within one week. For the experiment, the cleaning step in 2% Hellmanex was repeated twice and once with deionised water. These coverslips were used within 2 days. Prior to the hydrogel assembly, half of the coverslips were spin coated (1000 rpm, 1 min, WS-650Mz - 23NPPB Spin Coater, Laurell Technologies) with 20 *μ*l of a 1:250 dilution of multifluorescent beads (TetraSpeck^™^ beads, ThermoFisher, 0.1 *μ*m) allowing post-processing of the stage drift.

### Single-molecule staining for tracking single VSGs

1×10^7^ trypanosomes were harvested by centrifugation (10 min, 1400 xg, 4°C), followed by three washing cycles (1 min, 1400 xg, 4°C) of the cell pellet with 1ml TDB. After the resuspension in 100 *μ*l TDB, cells were chilled on melting ice (0°C) for 5 min to block endo- and exocytosis. Staining was performed at a final concentration of 2nM Atto-NHS 647N dye (ATTO-TEC) in a total volume of 200 *μ*l. Two washing steps with 1 ml TDB were subsequently carried out. Until the formation of the hydrogel, cells were kept in a volume of 1 ml TDB and at 0°C. For the embedding, the cells were resuspended in a volume of 10 - 20 *μ*l.

### Trypanosome immobilisation in hydrogels

Trypanosomes were efficiently immobilised in a hydrogel of 10% (w/v) vinylsulfone-functionalised polyethylenglycol (PEG-VS,tripentaerythritol core, Sigma-Aldrich) and 5.6% (w/v) thiol-functionalised hyaluronic acid (HA-SH, 24 kDa, ~ 54% SH-substituted, provided by group of Jürgen Groll) or 5% (w/v) thiol-functionalised hyaluronic acid (HA-SH, ~ 10 kDa, ~ 40% SH-substituted, provided by group of Jürgen Groll). Precursor solutions of PEG-VS (50% (w/v)) and HA-SH (24% (w/v)) were used for assembly of the hydrogel. Additives were spacer beads (non fluorescent FACS calibration, ∅ 6 *μ*m, BD Biosciences) and freely diffusing Atto-NHS 647N dye (ATTO-TEC) in a final concentration of 0.1nM in a total hydrogel volume of 10 *μ*l. This low concentration of initially unbound dye in the background may bind VSGs and replace bleached signals during imaging. This way, the label density was optimised. 2 - 3 *μ*l of concentrated trypanosome suspension was gently mixed with the components of the hydrogel. Polymerisation was started by the addition of HA-SH precursor solution. Immediately afterwards, 4 *μ*l of the hydrogel solution were placed between a bead-coated coverslip (bottom) and a non-coated coverslip (top). Subsequently, the top coverslip was weight down with a 90 mg weight, followed by centrifugation of the hydrogel sandwich in a centrifugation chamber to settle the cells in close proximity to the multifluorescent beads on the bottom coverslip (1500 xg, 1 min). The spacer beads prevented squeezing of the cells during this procedure. A scheme of an assembled hydrogel is shown in Fig 7.

**Fig 7.**
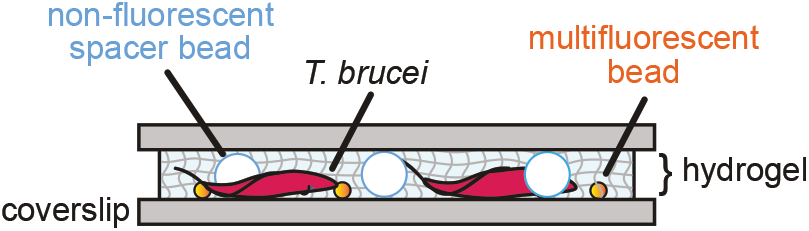
Scheme of an assembled hydrogel. It shows living trypanosomes immobilised in a hydrogel. The parasites were close to the bottom coverslip, which was spin-coated with multifluorescent beads used for drift correction. The addition of non-fluorescent spacer beads prevented squeezing during weighting in the assembling. Scheme is not to scale.

### Mechanical testing of the hydrogel

For the determination of the hydrogel stiffness, hydrogels with a polymer content of 10% (w/v) PEG-VS and 5% (w/v) HA-SH were cast in a cylindrical glass form (height = 4 mm, diameter = 6 mm). The measurements were performed directly after cross-linking using BOSE 5500 system (ElectroForce, Eden Prairie, MN, USA) for dynamical and mechanical testing. The total displacement was 2mm (50% compression) with a constant displacement rate of 0.005mm s^-1^. The Young’s Modulus was calculated from the raw data as the slope of the true stress-strain curve in the linear elastic range of 5 - 10% strain. The measurement was conducted in triplets.

### Single-molecule imaging and measurement

The single-molecule measurements were conducted at a setup based on an inverted-widefield microscope (Leica DMI6000B) equipped with a high numerical lens (HCX PL APO 100x 1.47 OIL CORR TIRF), a dichroic colour filter (zt405/514/633rpc, Chroma) and an EMCCD camera (iXon697, Andor technology). The region of interest was set to 120 x 120 px with a pixels size of 160 nm. Atto-NHS 647 *N* and eYFP molecules were sequentially excited with a 640nm laser beam at an intensity of 1 kW/cm^2^ and with a 514nm laser beam at an intensity of 2 kW/cm^2^. Signals were recorded in cropped-mode plus frame transfer and pulsed illumination of 9 ms. Each measurement comprised 10,000 - 20,000 consecutive images recorded at 100 Hz. The laser beams were shifted a few pixel horizontally relative to each other by the OptoSplit (CAIRN). Signal detection by the camera was enabled by a dichroic mirror (F48-635, Chroma) and the usage of the appropriate filter combination 550/49 BrightLine HC and 698/70 BrightLine HC (emission filters, Semrock) within the OptoSplit.

### Image processing

#### Localisation

Individual Atto-NHS 647N and eYFP molecules were localised with an average localisation precision of 26nm and 25 nm, respectively. Localisation was done using a MatLab routine (Mathworks Inc., USA) as described before in Schmidt et. al [15] and Fenz et. al [37]. In brief, a 2D-Gaussian fit was applied to the intensity profiles. The resulting list of molecule positions, width and intensities was filtered with respect to the known single-molecule footprints (width and intensity) as well as for detection error thresholds. The remaining localisations (x_i_, y_i_) and corresponding localisation precisions *σ*_i_ served as input for further data analysis.

#### Drift correction

Next to the cell of interest, movies also contained an immobile reference object (TetraSpeck^™^ bead, 100 nm) to correct the potential stage drift occurring during the measurements. The x- and y-positions of the reference object were smoothed with a built-in MatLab function smoothdata applying a Gaussian filter to dampen fluctuations in the order of the localisation precision. The filter size was set to 500 frames. The discrepancy of the smoothed positions to the starting position served as a reference for the correction of single-molecule positions in each frame.

#### Registration

A transformation matrix was generated from immobile, multifluorescent objects (TetraSpeck^™^ beads, 100 nm) that were acquired by alternating illumination in both colour channels. Preshifting of the green signals in an two pixel proximity to the red signals was performed prior to the generation of the transformation matrix with the MatLab built-in function images.geotrans.LocalWeightedMeanTransformation2D. The averaged error in the superposition precision was 55±5 nm.

#### Single-molecule tracking

The tracking algorithm has been described earlier in Fenz et. al [37]. In principle, trajectories were gained from correlation of the positional coordinates in successive images. The MatLab routine applied a probabilistic algorithm to identify all possible connections of all particles in neighbouring frames. From the generated transitional matrix the connection was chosen in the way that the total probability was maximised. The algorithm required an estimation of the diffusion coefficient as well as photophysical parameters to consider photobleaching and blinking behaviour.

### shortTrAn - Principles of calculations

In order to calculate the diffusion coefficient as well as the velocity from the displacements obtained by tracking VSGs, we used the method introduced by Hoze et. al [17], which is further described in Hoze and Holcman [12]. The evaluation of the displacements at the two distinct scenarios, diffusion and directed motion, was based on the Langevin model [38]. We adapted this method to our data and extended it in the aspects of the correction of the static and dynamic localisation error, the application of a spatial filter that takes the projection problem from 3*D* to 2*D* into account and the representation of the results from the calculation of the diffusion coefficient in an ellipse plot. This method allows the extraction of the local diffusion tensor and forces from a large number of short trajectories and will therefore be abbreviated as shortTrAn (short trajectory analysis).

#### Calculation of the local diffusion coefficient

A square with the side length *r* was centred at the position *X*. The one-dimensional local diffusion tensor *D^i,j^*(*X*, *r*) at position *X* was calculated from the mean square displacement of the total number of data points *N*(*X*, *r*) collected from trajectories *k* =1, 2, 3,…, *N* traversing the local square [12]:

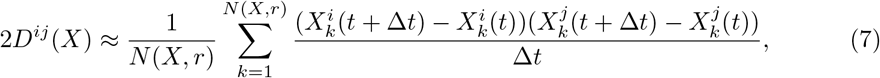

where *Deltat* was the time lag. Considering the static localisation error (*corr_stat_*) according to Martin et. al [19] and the dynamic localisation error (*corr_dyn_*) at non-continuous illumination according to Berglund [21] and Michalet and Berglund [22], in Eq 7 resulted in

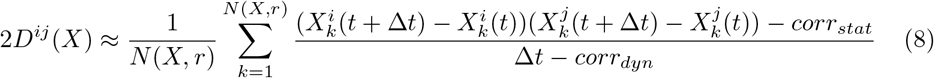

with *corr_stat_* = 2*σ*^2^ and 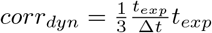. *σ* was the localisation precision and *t_exp_* was the exposure time of the camera. The two-dimensional average diffusion coefficient (*D*_(2*D*)_) was then calculated from the one-dimensional diffusion coefficients (*D^ii^*, *D^jj^*) by the following Eq 9:

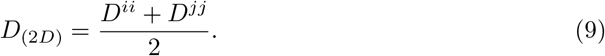

#### Calculation of the local directed motion

Simultaneously, the one-dimensional contribution of the local velocity *b_α_*(*X*, *r*) at position *X* was calculated by the approximation

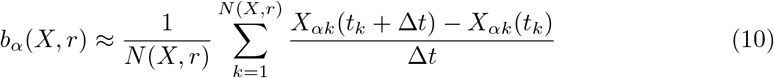

with *α* = *i*, *j* [12]. The two-dimensional velocity *b*(*X*, *r*) at position *X* was then determined by

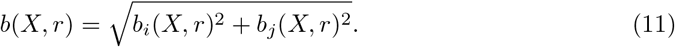

In order to consider the static localisation error in the local directed motion, the following assumption was made. The localisation precision of a point is defined with ±*σ* in one dimension. If the velocity of an object between two localisations is calculated, the total localisation error is ± 2*σ*. Subsequently, the relative standard error at the position *X* was determined by using the classical error propagation with *σ* = 26 nm. By dividing the relative standard error by 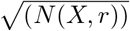, the local statistics were taken into account.

#### Calculation of the ellipse eccentricity

The eccentricity (*ε*) of an ellipse is defined by the following Eq 12:

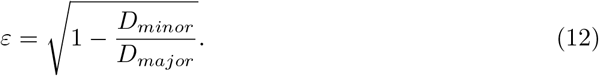

It is determined by the diffusion coefficient of the minor (*D_minor_*) and major axis (*D_major_*) of the ellipse, which was calculated from the tensor components by the usage of a transformation matrix (see “Implementation on empirical data in shortTrAn”).

### Implementation of shortTrAn on empirical data

A total number of 20 cells were analysed with the shortTrAn method. For each cell a large number of short trajectories served as an input and contained information about the trajectory identity, temporal succession, and the x/y-positions in pixel. First, the position units were transformed into nm with subsequent padding of a total 320 empty nm-spaces in each dimension to provide sufficient space for later binning.

To calculate the local diffusion coefficient and velocity, each trajectory was decomposed into its individual steps, the diffusion tensor and the velocity were calculated considering the static and dynamic localisation error and assigned to the location of the origin of the substep in the pre-defined multilayer feature space. There were three types of predefined feature spaces: The count field contained the number of localisations at position *i*, *j*. The tensor fields contained the results of the calculated tensor components *D_ii_*, *D_ij_*, *D_ji_* and *D_jj_*. The calculated one-dimensional velocities of the motion were placed in the corresponding vector fields *b_i_* and *b_j_*. The whole set of trajectories were iterated over. In the subsequent binning, the entries in the feature space were combined into larger units *S_i,j_* (binning size 160 nm) and then averaged over the number of contributing entries.

Three types of filters were then applied to the feature spaces: The first one removed entries from the function spaces which had low statistics. The second filter removed isolated squares. This served as a precautionary measure for the third filter, which removed completely isolated superpixels from the surface, as these did not contain any information about the VSG dynamics in the immediate vicinity. Finally, a filter combination was used to smooth out the data assuming similar conditions on the scale of neighbouring pixels (S3 Appendix) and to address the 3*D* to 2*D* projection problem at the rim of the trypanosome surface. After the filtering process, the diffusion coefficient and the velocity of an associated motion in two dimensions was then calculated by Eq 9 and Eq 11.

The diffusion tensors were visualised in an ellipse plot. The eigenvalues and eigenvectors were calculated from the tensor components using the transformation matrix ([cos,-sin;sin,cos]), where the eigenvalues reflect the diffusion value of the major and minor axes, while the eigenvectors define the angle of entry of the ellipse into the plot. The colour code represents the diffusion coefficient in 2*D*. Information about the velocity amplitude of the directed motion in 2*D* was encoded in a heat map, while the direction was represented in a quiver plot. The information about the static localisation error of the local directed motion was presented in a relative standard error map.

### Super-resolution image of the hook complex

The super-resolution image was computed as the sum of all fitted single - molecule positions. Each localisation (*x_i_*, *y_i_*) was plotted as a 2D Gaussian spot, where the respective width corresponded to the localisation precision *σ_i_* achieved for the eYFP molecule.

### Structural outline of the hook complex

The outline of the hook complex, which served as a reference for the entrance of the flagellar pocket, was generated by a self-written Python routine. The unit transformation to nm was applied to the x/y-position of the structural data, followed by a padding of a total 320 empty nm-spaces to provide sufficient space for later binning. Background and localisation outliers were manually removed and a concave hull of the remaining localisations was compiled with the Python built-in function alphashape.alphashape and an α parameter of 0.85. The coordinates of the hull vertices were scaled by the binning factor 160 to ensure compatibility with the shortTrAn created maps.

### Simulations

*In silico* experiments were carried out with the mean experimentally determined diffusion coefficient of 1.00 *μ*m^2^/s on trypanosome-shaped masks. The masks were a composite of a cellular outline and the corresponding flagellar pocket entrance region (FPR) marking the escape site. The cellular outline was determined from the count field, while the FPR was manually chosen from the part of the structural outline wrapped around the FP entrance at a resolution of 10 nm. To approximate the 3*D* shape of a trypanosome, the cellular outline layer was doubled excluding the FPR. 1,000 starting points were randomly distributed on the first layer. 2,000 dimensionless particles were simulated to perform a random walk on the mask until escape through the FPR. When interacting with the boundaries of the current layer, the particle was relocated to the other layer. Simulation settings are illustrated in Fig 8. For each step a random angle was chosen *α* ∈ [0, 2 *π*] while the step lengths (*l*) fulfilled the criterion for the probability (*p*) in Eq 13 [37]:

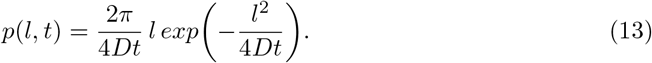

where *D* was the diffusion coefficient and *t* the time lag. The individual escape times (*ET*) from all particles were collected and plotted in a probability density histogram. The mean first passage time (*τ*) was determined from the fit in the exponential tail of the histogram:

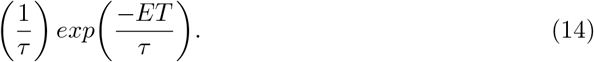

The tail region was found automatically by minimising the corresponding *R*^2^ value. We chose the trajectory length long enough to ensure almost all particles were able to escape during simulation (50,000 steps corresponding to 500 s).

**Fig 8.**
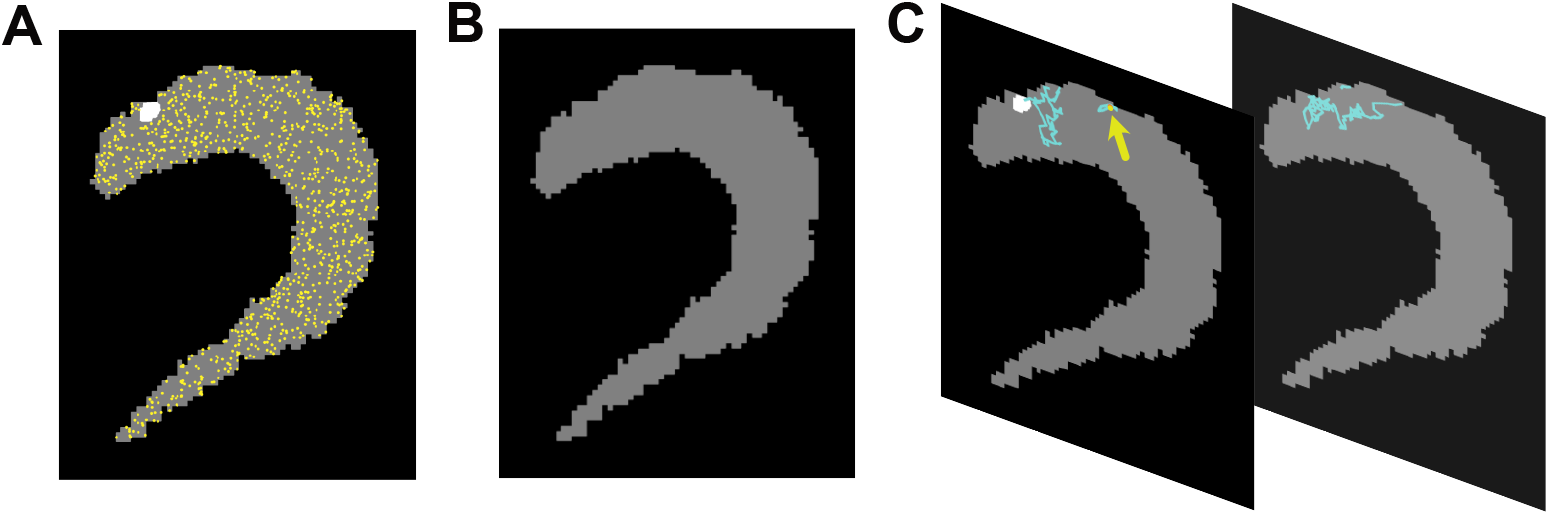
Random walk simulation on trypanosome-shaped masks. **(A)** First leaflet (dark grey) contained the flagellar pocket entrance region (white) and the starting positions (yellow). **(B)** The second leaflet (dark grey) extends the area to mimic the 3*D* plasticity of a trypanosome. It contained no starting points and no flagellar pocket entrance region. **(C)** Representation of a simulated random walk trajectory (cyan). Only the last 2 seconds are shown for clarity. The starting point is marked by a yellow arrow. The trajectory switched three times the leaflet until it escaped by entering the flagellar pocket entrance region.

## Data availability

The shortTrAn algorithm is available at https://github.com/MarSc13/shortTrAn as Python scripts and Jupyter Notebooks. The repository contains the source code as well as exemplary data from the measurements. Further data can be provided upon request. We have also used Zenodo to assign a DOI to the repository: 10.5281/zenodo.6941359.

## Supporting information

**S1 Fig.**
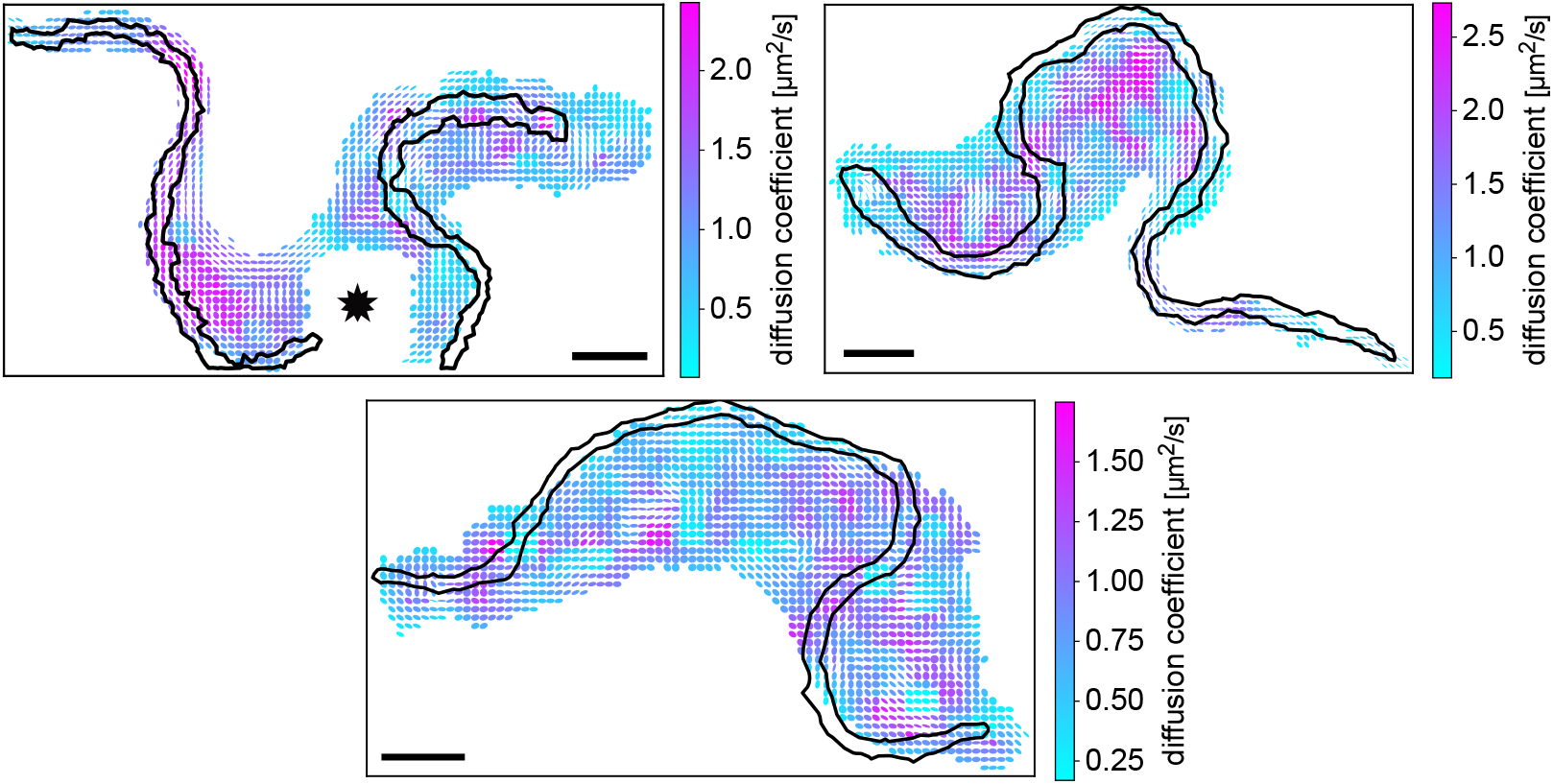
Local diffusion coefficients in relation to the flagellum. Diffusion maps of three trypanosome surfaces are displayed, with the 2*D* diffusion coefficient colour-coded. The shape of the ellipses indicates isotropic (circle) or anisotropic (ellipse) diffusion. The black line represents the outline of the flagellum on the pellicular membrane. The area outshone by a TS bead is marked by an asterisk. The scale bars are 1.6 *μ*m.

**S2 Fig.**
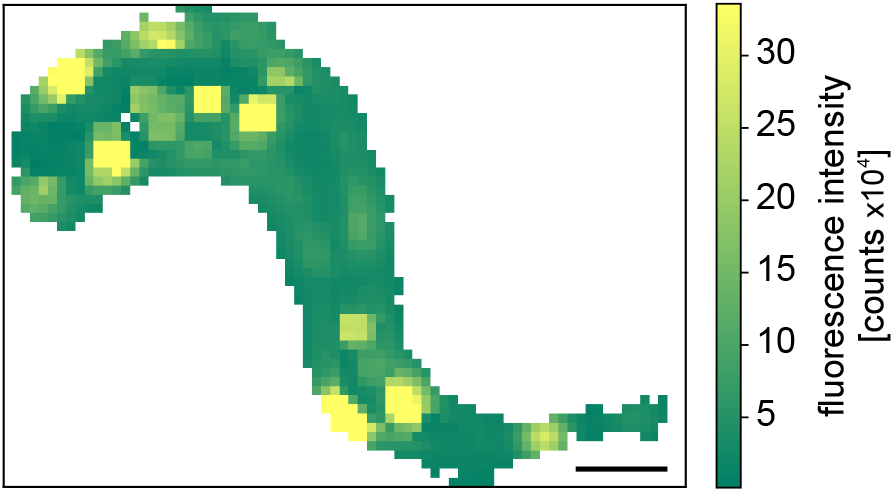
Cumulative VSG fluorescence encoding the dwelling time. Cumulative VSG fluorescence encoding the dwelling time. The heat map displays the sum of all emitters’ fluorescence detected within each superpixel of the exemplary cell. The scale bar is 1.6 *μ*m.

**S3 Fig.**
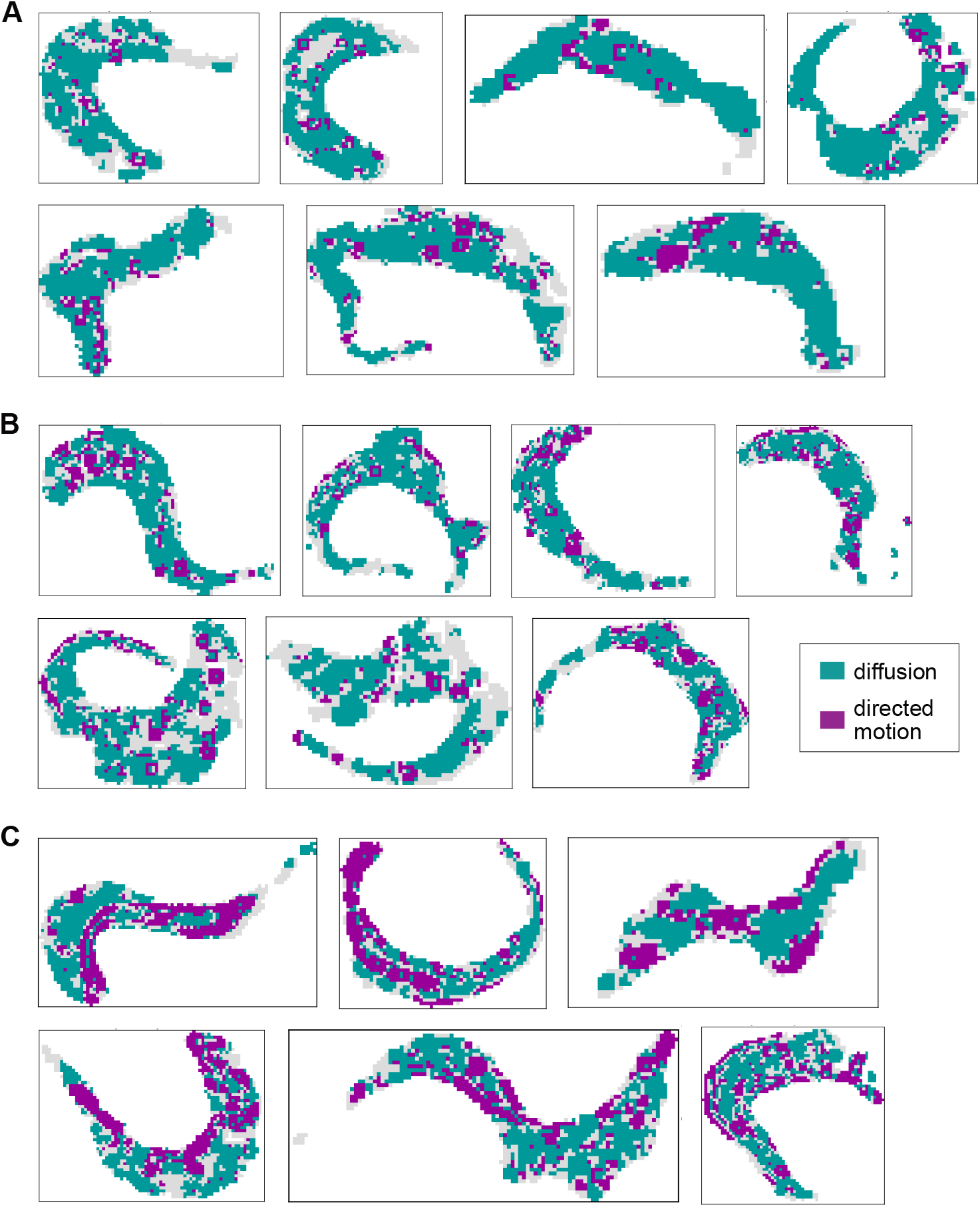
Compilation of all supermaps generated from tracking VSGs on bloodstream trypanosomes. Superpixels in green represent diffusion, in magenta directed motion and grey are superpixels that were not explicitly assigned to one of the two motion models. Each superpixel is of the size 0.16 x 0.16 *μ*m. **(A)** Group 1 was characterised by mainly diffusive superpixels (*N* = 7 supermaps). **(B)** Group 2 comprised a higher proportion of superpixels assigned to directed motion, which were arranged in patches (*N* = 7 supermaps). **(C)** Supermaps categorised to group 3 were featured by a large number of superpixels assigned to a directed motion, which were additionally arranged in elongated traps (*N* = 6 supermaps).

## S1 Appendix. Verification of the established eYFP::MORN1 cell line

In order to analyse the surface VSGs dynamics in relation to the flagellar pocket a 13-90 cell line was established expressing an N-terminally eYFP tagged TbMORN1 (MORN1). To facilitate the localisation of single molecules, the tagging was performed of only one allele. MORN1 is one component of the hook complex and distributed all over the full structure [13, 14]. To verify that the fusion protein is expressed and behaved like the WT protein, the cell growth rate was observed, western blot analysis were performed (shown in Fig S1-1) and the structure of the hook complex was resolved by high-resolution microscopy.

**Fig S1-1.**
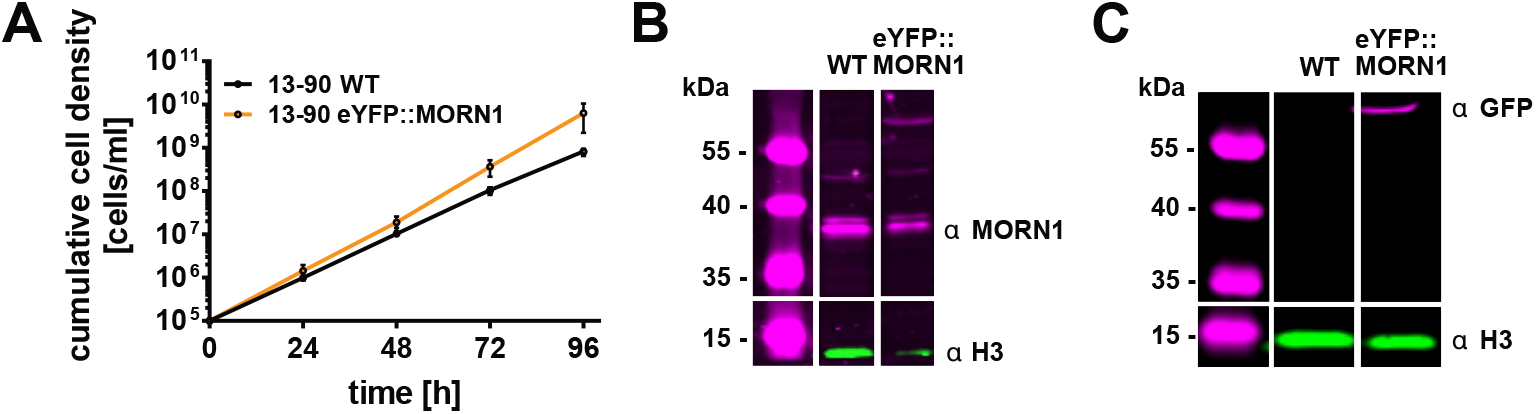
Cell viability of the 13-90 eYFP::MORN1 cell line. **(A)** Cumulative growth curve of the parental cell line 13-90WT and the transgenic 13-90 eYFP::MORN1 cell line. **(B)** Western Blot analysis with an antibody against MORN1 confirmed the expression of the WT and endogenously eYFP-tagged allele. The detection of the histone variant H3 was used as loading control. **(C)** Detection of the expression of the endogenously eYFP-tagged MORN1 with an GFP-antibody in Western Blot analysis. The histone variant H3 served as loading control.

The observation of the growth rate of the transgenic cell line in comparison to its parental cell line over 72 h exhibited no growth defect (Fig S1-1 A) with similar doubling times for the WT (7.42±0.08 h) and transgenic cell line (6.24±0.42 h). Western blot analysis revealed the expression of the MORN1 tagged with eYFP in addition to the WT protein. The antibody against MORN1 detected the MORN1 WT protein at 40 kDa as well as the fusion protein at 67 kDa (Fig S1-1B). The antibody against GFP could clearly confirm the expression of the fusion protein (67 kDa, Fig S1-1 C). This, together with the resolution of a hook-like structure of the hook complex with single-molecule fluorescence microscopy (Fig 1 B), indicated that the fusion protein was behaving as expected.

## S2 Appendix. Determination of the single-molecule localisation precision

We assessed the localisation precision *σ* by three different methods: (i) From the error of the x- and y-position in the Gaussian fit, (ii) from theory, and (iii) from the standard deviation in the x- and y-position of an immobile emitter. The results from the different methods are shown in the following Table S2-1, while the details of the methods are described in the following sections.

**Table S2-1.** Obtained localisation precisions (*σ*) from the error in the Gauss fit of the single-molecule detection, from theory and from the standard deviation in the x- and y-position of an immobile emitter. The final *σ* was calculated from the mean value.

| | $\sigma$ |
| --- | --- |
| Gauss fit | 25 nm |
| Theory | 28 nm |
| Immobile particle | 26 nm |
| Average | 26 nm |

### (i) Determination from the Gauss fit

When localising the single molecules, the localisation precision *σ* can be derived from the error of the x- and y-position determined by the fit of a 2*D* Gaussian to the intensity profile of the single molecule. In our case, an average was extracted from the error over all localisations of the 20 different cell datasets. The mean value was 25 nm.

### (ii) Determination from theory

The localisation precision is defined in theory [23] according to the following equation:

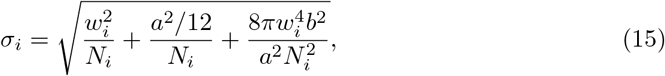

where *w_i_* is the width of the Gaussian, *N_i_* is the number of emitted photons, *a* is the pixel size and *b* is the background noise. The number of incoming photons for each localisation (*N_i_*) was derived from the counts (*C*) registered on the chip of the EMCCD with the help of the count conversion guide provided by Andor, which is displayed below:

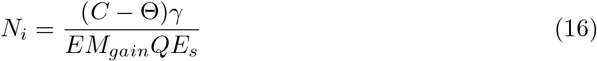

with the offset of the background *θ*, the camera sensitivity *γ* of 10.7 electrons per A/D unit, the EM gain (*EM_gain_*) of 150 and the sensor quantum efficiency (*QE_s_*) of 0.94, which was extracted from camera specification sheet for the emission wavelength 664nm of the Atto-NHS 647N dye. The offset of the background *θ* was negligible because a flat field correction was applied during the measurement to remove the background signal detected in the dark. For the determination of the theoretically *σ*, localisations of eleven immobile Atto-NHS 647N emitters were extracted with a minimum of 100 localisations. The theoretically calculated localisation precision *σ* was 28 nm. See the next section for a protocol on how to prepare immobile emitters.

### (iii) Determination from the standard deviation of an immobile emitter

In the third method, *σ* can be determined from the standard deviation of the x- and y-position of an immobile emitter. In order to obtain immobile emitters, a methanol fixation protocol was chosen, as it was shown that membrane anchored proteins are inefficiently immobilised with formaldehyde [39]. Trypanosomes were labelled with a nanomolar quantity of Atto dye according to the method section and fixed in −20°C methanol for 15 min at −20°C. They were then washed three times with 1 ml TDB (1400 xg, 1 min, 4°C). Thereafter, the cells were embedded in the hydrogel for microscopy. To guarantee that a sufficient number of localisations sample the range of the localisation precision, only trajectories with a minimum length of 100 steps were selected. Eleven emitters were analysed. The reported value of *σ* = 26nm was the smallest value found. The corresponding trajectory had a length of 480 steps and a round shape. We decided on the smallest value because in this case the emitter was most likely completely immobilised while in the cases with a larger area covered by the particle trace residual movement could not be ruled out.

## S3 Appendix. Spatial filters

### First spatial filter

In order to remove pixels with low statistics from the analysis, a threshold was applied to the count field *cf* which resulted in binerisation and set the superpixel *S* at position *i*, *j* with a population number *N* of ≤ 5 to zero (see Eq 17). Multiplication with the count field itself, the tensor fields and the vector fields removed the entries in the corresponding superpixel *S_i,j_*.

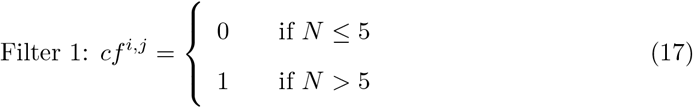

### Second spatial filter

It was the purpose of the second spatial filter to remove isolated superpixels from the analysis. This served as a precautionary measure for the third spatial filter as the immediate environment did not contain any information to allow for smoothing out the data. To this end, the binarised count field served as input to Eq 18 as follows:

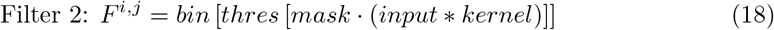

Convolution of the binarised count field with a box kernel of size 3 superpixels resulted in the number of populated superpixels of the adjacent neighbourhood (max. connectivity of 8). Multiplication of the convoluted matrix with the input as a mask resulted in the elimination of superpixel which were filled even though they were not originally populated. Application of a threshold of ≤ 1 removed isolated superpixel. The final binerisation, which masked empty superpixel with 0 and populated superpixel with 1, yielded the filter matrix *F^i,j^*. This matrix was then applied to the count field itself, tensor and vector matrices for the removal of entries in isolated superpixel. The workflow describing Eq 18 of the second spatial filter is depicted in Fig S3-1.

### Third spatial filter

For smoothing out the data, we used a sliding box kernel of the size three superpixels simultaneously on the count field *cf* and input field *if*, either a tensor or vector field, to calculate the weighted average at position *i*, *j* (Eq 19).

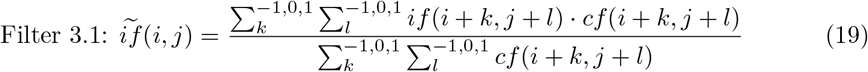

The 3*D* to 2*D* projection issue at the rim of the trypanosomes was accounted by applying Filter 3.2 which used the Python built-in Gaussian scipy.ndimage.gaussian_filter [40] on the duplicated matrix already applied with Filter 3.1. The Gaussian filter was of a radius of 6 superpixels, with a standard deviation *σ* for the Gaussian kernel of 1 and constant mode with a value of zero. This means that the superpixels outside the trypanosome mask needed for the Gaussian were populated with the value 0. Finally, the results from the smoothing and the projection filter were combined in the following way: Superpixels at the rim were populated with the results of the projection Filter 3.2, while superpixels of the surface area were populated with the results of the smoothing Filter 3.1 (see Fig S3-2).

**Fig S3-1.**
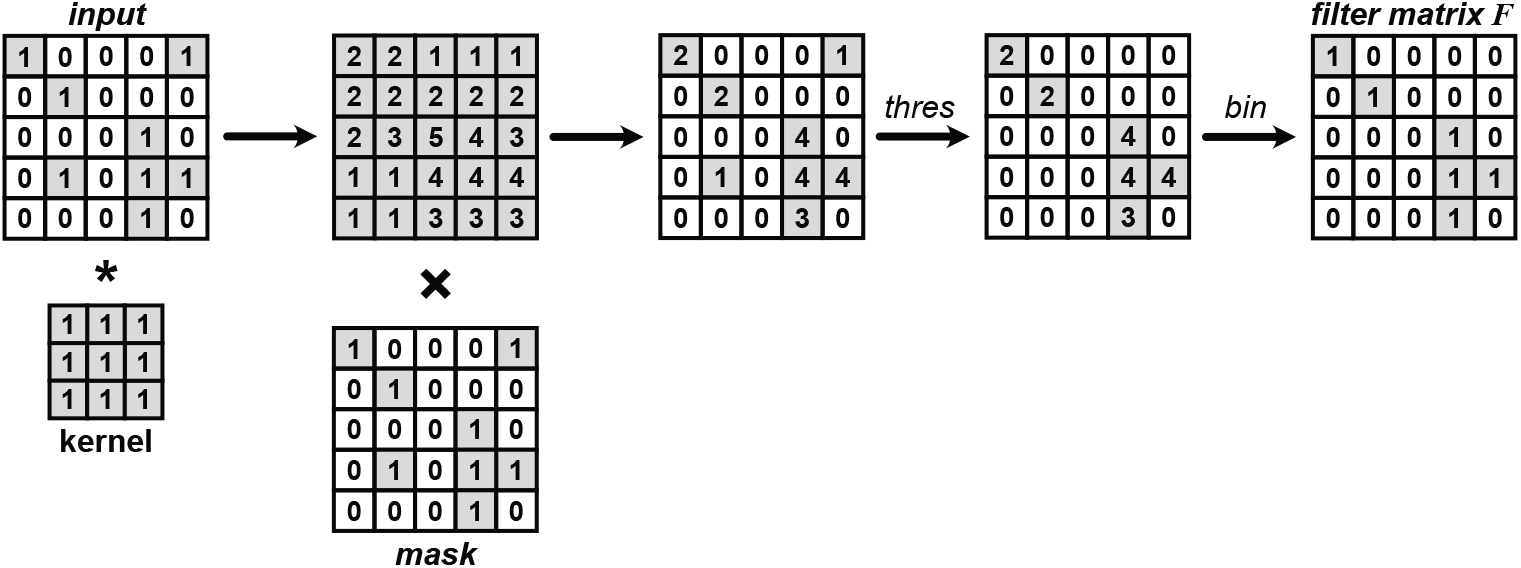
Generation of the *filter matrix F* for the second spatial filter. Convolution of the binarised count field with a box kernel of the size three superpixel yielded a matrix containing the number of populated superpixels of the directly adjacent neighbourhood. The removal of falsely positive populated superpixels was accomplished by the multiplication of the binarised count field (*mask*). Application of a threshold of one eliminated isolated superpixel. The final *filter matrix F* was obtained by binerisation. Application to the count field, tensor and vector matrices enabled the removal of isolated superpixels.

**Fig S3-2.**
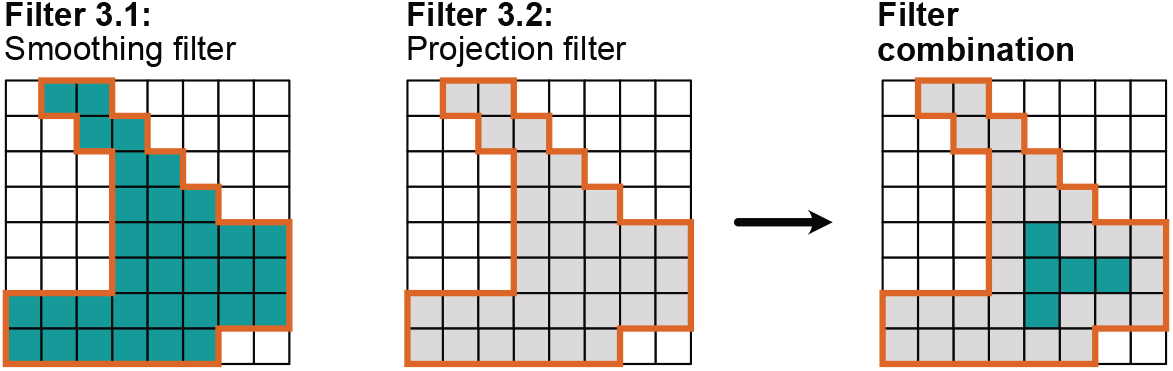
Combination scheme of the third spatial filters. The matrices obtained from the application of Filter 3.1 and Filter 3.2 served as input. The rim of the surface area was determined and used to populate superpixels at the rim with the results of the Filter 3.2, while the centre was populated with the results of Filter 3.1.

## S4 Appendix. Necessity of a projection filter for measurement results at the trypanosome rim

The implementation of the algorithm on the VSG tracking data indicated an accumulation of high velocities especially at the rim of the cell surface (shown in FigS4-1) and thus the need to introduce a 3*D* to 2*D* projection filter in the spatial filters.

**Fig S4-1.**
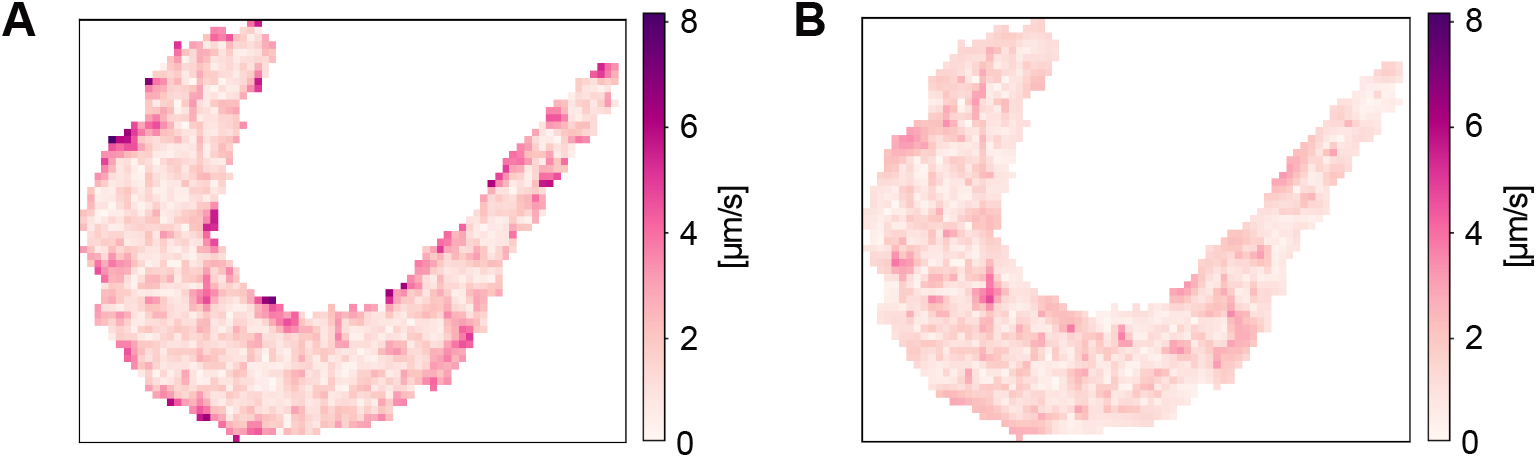
Artefacts of high velocities at the trypanosome rim. **(A)** Heat map of random walk simulation reveals artefacts of high velocities if the smoothing filter is applied solely. **(B)** Heat map of the same simulation after the additional application of the projection to the smoothing filter.

This arose because the results at a superpixel are an average value over this area. If the superpixel was in the centre of the cell surface, movements in all directions in the 2*D* plane from this superpixel could be registered, resulting in a circle in the ellipse plot in the diffusion representation. The calculated velocities of the directed motion at the location with diffusion would cancel each other out in two dimensions due to the uniform contribution to all directions. If the region of interest was directly at the rim of the cell surface, movements following the 3*D* shape and thus out of the focal plane could not be accounted for. As a consequence, free diffusion appeared anisotropic whereas in the directed motion scenario, the result was an unusually large velocity in direction perpendicular to the rim. The underlying principle is visualised in Fig S4-2. The width of the filter was validated by random walk simulations on the trypanosome mask evaluated with shortTrAn. To correct the high velocities, we have therefore implemented a Gaussian filter in the third spatial filter, which is placed over the outer two superpixels. The width of the filter was adjusted until the velocities of the superpixels at the rim were found to be in the same range as the velocities in the remaining superpixels. The implementation of the third spatial filter is more precisely elucidated in S3 Appendix.

**Fig S4-2.**
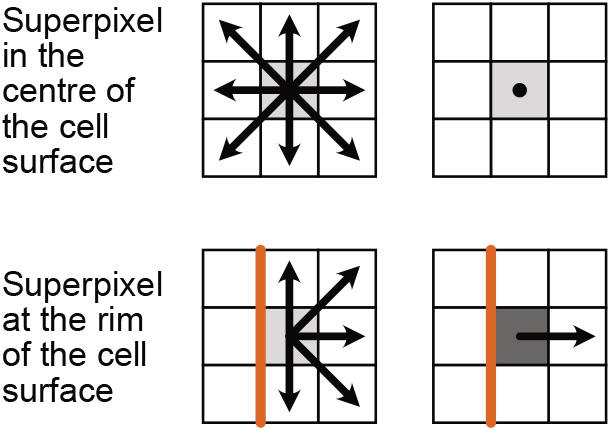
Principle for the emergence of the projection error. Entries of a superpixel are an averaged value from single-step events emanating from this superpixel. Under the assumption of diffusion, velocities within a superpixel would cancel each other out due to the uniform contribution to all directions. It is clearly visible at superpixels located in the centre of a cell surface projection. If the superpixel of interest is located at the rim of the cell surface, movements following the 3*D* shape are out-of-focus and could not be accounted for. Consequently, superpixels show a large net effect of velocities in direction perpendicular to the rim. The velocity amplitude is depicted in different shades of grey. The darker, the higher is the value.

## Acknowledgments

We are grateful to Thomas Schmidt for providing MATLAB code to localise single-molecules and perform single-molecule tracking. Furthermore, we would like to thank David Holcmann and Matthias Weiss for the helpful discussions on the implementation and Christian Janzen for providing an antibody for WB analysis.

## Author Contributions

Conceptualisation, M.S., T.P., M.G., M.E., P.K., S.F.; Methodology, M.S., M.G., L.F., B.M., J.T., J.G., S.F.; Software, M.S., T.P., M.G., P.K., S.F.; Formal Analysis, M.S., T.P., P.K., S.F.; Investigation, M.S., L.F.; Writing - Original Draft, M.S., S.F.; Writing - Review & Editing, all authors; Supervision, J.T., J.G., M.E., P.K., S.F.; Funding Acquisition, J.T., J.G., M.E., S.F.

